# The Arabidopsis leucine rich repeat receptor-like kinase MIK2 interacts with RKS1 and participates to the control of quantitative disease resistance to the bacterial pathogen *Xanthomonas campestris*

**DOI:** 10.1101/2024.01.29.577741

**Authors:** Florent Delplace, Carine Huard-Chauveau, Fabrice Roux, Dominique Roby

**Affiliations:** Laboratoire des Interactions Plantes-Microbes Environnement (LIPME), INRAE, CNRS, Université de Toulouse, 31326 Castanet-Tolosan, France

**Keywords:** plant-pathogen interactions, immunity, quantitative disease resistance, receptor, protein-protein interaction

## Abstract

Molecular mechanisms underlying qualitative resistance have been intensively studied. In contrast, although quantitative disease resistance (QDR) is a common, durable and broad-spectrum form of immune responses in plants, only a few related functional analyses have been reported. In this context, the atypical kinase RKS1 is a major actor of QDR to the bacterial pathogen *Xanthomonas campestris* (*Xcc*) and is positioned in a robust protein-protein decentralized network. Among the putative interactors of RKS1 found by yeast two hybrid screening, we identified the receptor like kinase MDIS1-Interacting Receptor-like Kinase 2 (MIK2). Here, by multiple and complementary strategies including protein-protein interaction tests, mutant analysis and network reconstruction, we report that *MIK2* is a component of *RKS1* mediated QDR to *Xcc*. First, by co-localization experiment, co-immunoprecipitation (Co-IP) and Bimolecular Fluorescence Complementation (BiFC), we validated the physical interaction between RKS1 and MIK2 in the plasma membrane. Using *mik2* mutants, we then showed that *MIK2* is required for QDR at the same level as *RKS1*. Interestingly, a catalytic mutant of MIK2 was able to interact with RKS1 but unable to fully complement the *mik2-1* mutant in response to *Xcc*. Finally, we investigated a potential role of the MIK2-RKS1 complex as a scaffolding component for coordination of perception events, by constructing a RKS1-MIK2 centered protein-protein network. Eight mutants corresponding to seven RLKs of this network showed a strong and significant alteration in QDR to *Xcc*. Our findings provide new insights into the molecular mechanisms underlying perception events involved in QDR to *Xcc*.

## Introduction

In response to pathogens, plants have developed complex resistance mechanisms that are either constitutively expressed or induced after a pathogen attack (Glazebrook, 2005; Panstruga *et al*., 2009). Quantitative disease resistance (QDR) is the predominant form of resistance in crops and natural populations. QDR is characterized by a polygenic determinism conferring partial and generally broad-spectrum resistance (Poland *et al*., 2009; Roux, *et al*., 2014b). For this reason, the molecular mechanisms underlying QDR remain poorly understood. However, the recent cloning of a limited number of QDR genes underlying resistance QTLs revealed a broad range of molecular functions (Delplace *et al*., 2021). Only a very few QDR genes encode NLRs (nucleotide-binding leucine-rich repeat receptors) identified as *R* genes in the context of qualitative resistance (Broglie *et al*., 2006; Staal *et al*., 2006, Fukuoka *et al*., 2014; Demirjian *et al*., 2023). On the other hand, several QDR genes encode different types of receptors including receptor-like kinases (RLKs), signaling components such as kinases, and diverse metabolic functions (Corwin and Kliebenstein, 2017; Pilet-Nayel *et al*., 2017; Nelson *et al*., 2018). This diversity of molecular functions together with the partial resistance conferred by QDR genes suggest that QDR results from a complex network integrating diverse pathways in response to multiple pathogenic determinants (Roux *et al*., 2014b). This suggests in turn that (i) individual QDR genes may confer resistance to multiple pathogens (Nelson *et al*., 2018; Kanyuka and Rudd, 2019), and (ii) multiple receptors act together to perceive diverse pathogen determinants (Pok Man Ngou *et al.,* 2022). However, a limited number of receptors have been identified in the context of QDR in various plant-pathogen interactions.

In plants, receptor kinases (RKs) play an essential role in environmental signal perception, including pathogen detection at early stages of infection. RKs detect different pathogen epitopes. The transmembrane RK Flagellin Sensitive 2 (FLS2) is implicated in the detection of flg22, a conserved peptide of bacteria flagellin (Gómez-Gómez and Boller, 2000; Bauer *et al*., 2001). The EF-Tu receptor (EFR) is involved in the perception of the peptide elf18, the N-terminal peptide of EF-Tu (Kunze *et al*., 2004; Zipfel *et al*., 2006). However, the number of distinct perception systems with specificity for microbe-associated molecular patterns (MAMPs) is difficult to estimate and a limited number of RKs have been described to recognize pathogen signatures (Tang *et al*., 2017). Recently, a complex and dynamic network of LRR-RKs interactions has been described in *Arabidopsis thaliana* (Smakowska-Luzan *et al*., 2018). The association of LRR-RKs in heterodimers might participate to a large detection of diverse epitopes involved in various plant processes. RK dependent recognition events converge into receptor hubs and overlapping immune signaling components can be recruited (Holton *et al*., 2015; Adachi and Tsuda, 2019). These RKs take place in plasma membrane protein complexes including other RLKs, receptor proteins (RPs), kinases or pseudokinases (Monaghan and Zipfel, 2012; Boudeau *et al*., 2006) and play a central role in the modulation of plant immunity. Some pathogen effectors have been shown to physically interact with essential signaling components to inhibit fast information spreading (Ahmed *et al*., 2018). In the same vein, complex associations between NLRs have been observed mainly in heterodimers (Wróblewski *et al*., 2018), and most cell surface and intracellular immune receptors appear to engage other receptors including ligand-binding receptors and transducer co-receptors (Kamoun *et al*., 2018). In summary, pathogen detection is not limited to a ligand perception event and receptors are organized in complex networks involving multiple perception events, in order to recognize a large variety of pathogen determinants. However, only a few of such ligand-binding receptors or receptor complexes have been identified and reported in the literature.

In the plant - pathogen interaction between *A. thaliana* and the vascular bacterium *Xanthomonas campestris* pv. *campestris* (*Xcc*), RKS1 (Resistance related KinaSe1) was identified to confer QDR to *Xcc* (Huard-Chauveau *et al*., 2013). RKS1 encodes an atypical kinase lacking some critical domains in the kinase catalytic core (Huard-Chauveau *et al*., 2013; Roux *et al*., 2014a). Atypical kinases (or pseudokinases) have been described as important regulators of signaling networks (Reiterer *et al*., 2014; Blaum *et al*., 2014). Recently, we reported a highly interconnected and decentralized RKS1-dependent protein-protein network, which is largely distinct from Effector-Triggered Immunity (ETI) and Pattern-Triggered Immunity (PTI) responses already characterized in *A. thaliana* (Delplace *et al*., 2020). From this network, we identified a signaling subnetwork that includes RLKs not previously described in the context of plant immunity for most of them. Thus, functional analysis of RLKs associated with RKS1 could provide a powerful tool for deciphering molecular mechanisms involved in the early signaling events of QDR. In this study, we report that the LRR-RLK MDIS1-Interacting receptor-like Kinase 2 (MIK2) physically interacts with RKS1 and that MIK2 and RKS1 co-localized at the plasma membrane. We also show that *mik2* mutants exhibit increased susceptibility to *Xcc* as compared to the wild type and that MIK2 catalytic activity is required for QDR to *Xcc*. In addition, the double mutant *mik2-1/rks1-1* showed a similar level of resistance against *Xcc568* as the simple mutants, indicating that they probably belong to the same pathway. Interestingly, MIK2 was recently demonstrated as a ligand-perceiving RLK (Hou *et al*., 2021; Rhodes *et al*., 2021). Accordingly, we found that MIK2 is positioned in a putative receptors-interacting network. Insertional mutants for some of these RLKs showed increased susceptibility to *Xcc,* suggesting that they participate with MIK2 to *Xcc* perception. Our data support the hypothesis that MIK2 and RKS1 act as a scaffolding hub in a RLK network to centralize and modulate pathogen determinant perception for QDR signaling and activation.

## Results

### RKS1 physically interacts with MIK2, a leucine rich repeat receptor-like kinase

Using a Yeast two hybrid screen and a mutated version of RKS1 (RKS1^D191A^) as a bait against a cDNA library generated from leaves inoculated by the strain *Xcc147* (Froidure *et al*., 2010), we identified among 43 candidate proteins (Delplace *et al*., 2020), a fragment of the kinase domain of MIK2 as a putative interactor of RKS1. We investigated the interaction between RKS1 and MIK2 using several strategies. Firstly, to test whether RKS1 and MIK2 might be localized in the same cellular compartment, constructs using the fluorescent C-terminal tags eGFP and mRFP1 respectively fused to the full-length sequences of RKS1^D191A^ and MIK2, were co-transfected into Arabidopsis seedlings. As previously reported (Delplace *et al*., 2020), RKS1^D191A^-eGFP was detected in the plasma membrane, the cytoplasm and the nucleus. MIK2 was detected only in the plasma membrane (Fig. 1A). An overlap between eGFP and mRFP1 signals indicates a close subcellular localization of MIK2 and RKS1 in the plasma membrane. Secondly, co-immunoprecipitation assays were performed using RKS1^D191A^ fused to a c-myc tag and the kinase domain of MIK2 (MIK2-KD) fused to aHA (Hemagglutinin) tag. After immunoprecipitation with an anti-HA antibody, RKS1^D191A^-c-myc was detected when incubated with MIK2-KD-HA (Fig. 1B). ZAR1-HA, used as a positive control of interaction with RKS1 (Wang *et al*., 2015) and RKS1^D191A^-c-myc, was not detected after immunoprecipitation with tag-HA, indicating the specific nature of interaction between MIK2 and RKS1 (Fig. 1C). An immunoprecipitation of RKS1^D191A^-HA with an anti-HA antibody was also able to pull down MIK2 full-length protein with a c-myc tag (Supplemental Figure S1). Thirdly, we performed Bimolecular Fluorescence Complementation (BiFC) assays using the split YFP system. The N-terminal half of the yellow fluorescent protein was fused to *RKS1* and the YFP C-terminal was fused to *MIK2*, *ZAR1* or only the HA tag. When *RKS1-nYFP* was co-transfected with *MIK2-cYFP*, YFP signals were observed in the plasma membrane in *A. thaliana* protoplasts. As a comparison, when *RKS1-nYFP* was co-expressed with *ZAR1-cYFP*, YFP signals were detected in the plasma membrane and the cytoplasm (Fig. 1D). No YFP signal was observed with RKS1-nYFP and HA-cYFP co-transfected into *A. thaliana* protoplasts. These results collectively demonstrate that RKS1 is able to physically interact with the kinase domain of MIK2 in the plasma membrane.

**Figure 1.**
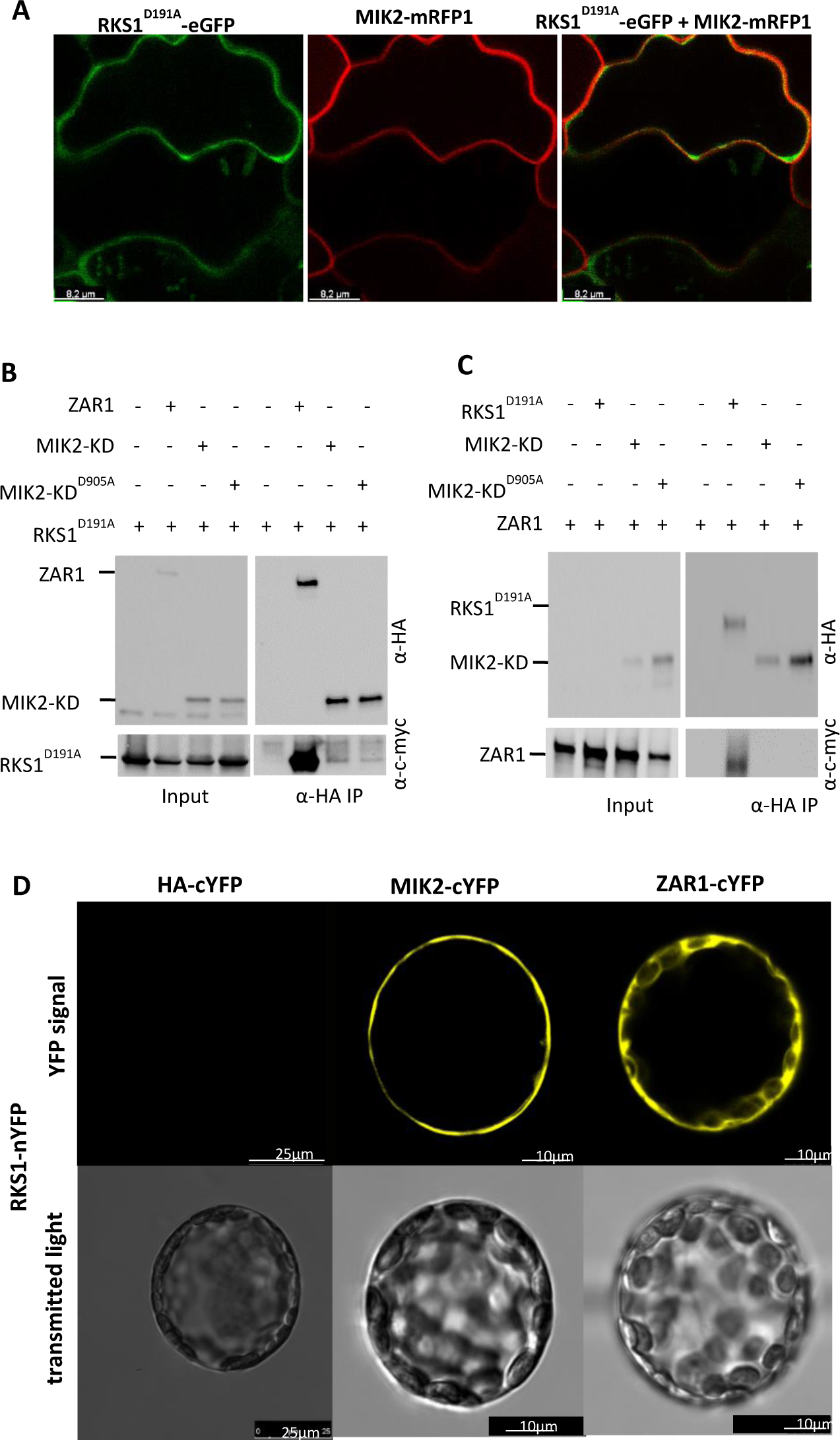
MIK2 physically interacts with RKS1 in the plasma membrane but not with ZAR1. **A,** Arabidopsis plants were co-transformed by RKS1^D191A^-eGFP and MIK2-mRFP1 full length constructs. Green corresponds to GFP signal, red corresponds to RFP signal and orange, co-localization between GFP and RFP. The white bars represent the ladder scale. **B,** Co-Immunoprecipitation experiments with protein extracts from *N. benthamiana* inoculated with the C58C1 *Agrobacterium* strain carrying *ZAR1-HA*, *MIK2-KD-HA* or *MIK2-KD^D905A^*-HA constructs, and with the C58C1 strain carrying *RKS1^D191A^-c-myc*. Input proteins were extracted and revealed by anti-HA and anti-c-myc HRP immunoblotting. Then, total proteins were subjected to anti-HA immunoprecipitation (α-HA IP), separated on a 4-15% SDS-PAGE gel and detected by anti-HA and anti-c-myc immunoblotting.**C,** Co-Immunoprecipitation experiments with protein extracts from *N. benthamiana* inoculated by the C58C1 *Agrobacterium* strain carrying RKS1^D191A^-HA, MIK2-KD-HA or MIK2-KD^D905A^-HA constructs in one hand, and in the other hand, with a strain carrying ZAR1-c-myc. Proteins were analyzed by immunoblotting as described in part B. **D,** Bimolecular Fluorescence Complementation (BiFC) assay between RKS1 and MIK2, ZAR1 or HA. RKS1 was fused to nYFP and MIK2, ZAR1 and HA were fused to cYFP. The indicated constructs were co-transformed into Arabidopsis protoplasts. The top panel represents the YFP signal and the bottom panel, the transmitted light. Bars in bottom right corner indicate the scale (25 or 10 μm).

### *MIK2* partially controls *RKS1*-dependent QDR

To investigate the function of *MIK2* in *Xcc* resistance, we characterized two T-DNA homozygous mutant lines for *MIK2*, *i.e. mik2-1* and *mik2-2* (SALK collection, Col-0 background). The flanking regions of the T-DNA insertion sites were sequenced and the T-DNA insertion sites were found in the first exon of *MIK2*, at +2015 bp and +2644 bp respectively (Fig. 2A). Complementarily, we generated transgenic lines complemented by *AtMIK2* in the *mik2-1* mutant background. *MIK2* gene expression in *mik2* mutants and *mik2-1* complemented lines is presented in Supplemental Fig. S2. While 31% and 5% of residual *MIK2* gene expression were found in *mik2-1* and *mik2-2* mutants respectively, *MIK2* gene expression was restored in the complemented *mik2-1(35S::MIK2) #1* and *#2* lines (629% and 57%), as compared to Col-0. *mik2* mutants and *mik2-1* complemented lines were then inoculated with a bacterial suspension of the strain *Xcc568* and disease index evaluated at 3-, 5-, 7-and 10-days post inoculation (Fig. 2B and 2C). *mik2* mutants showed a significant increased susceptibility compared to Col-0, and this increased susceptible phenotype was not statistically different from the *rks1-1* mutant phenotype. These results were confirmed by measurement of *Xcc*568 bacterial growth *in planta* (Supplemental Fig. S3). Complemented lines showed a similar resistance level than Col-0. These data confirm the implication of *MIK2* in QDR to *Xcc568*.

**Figure 2.**
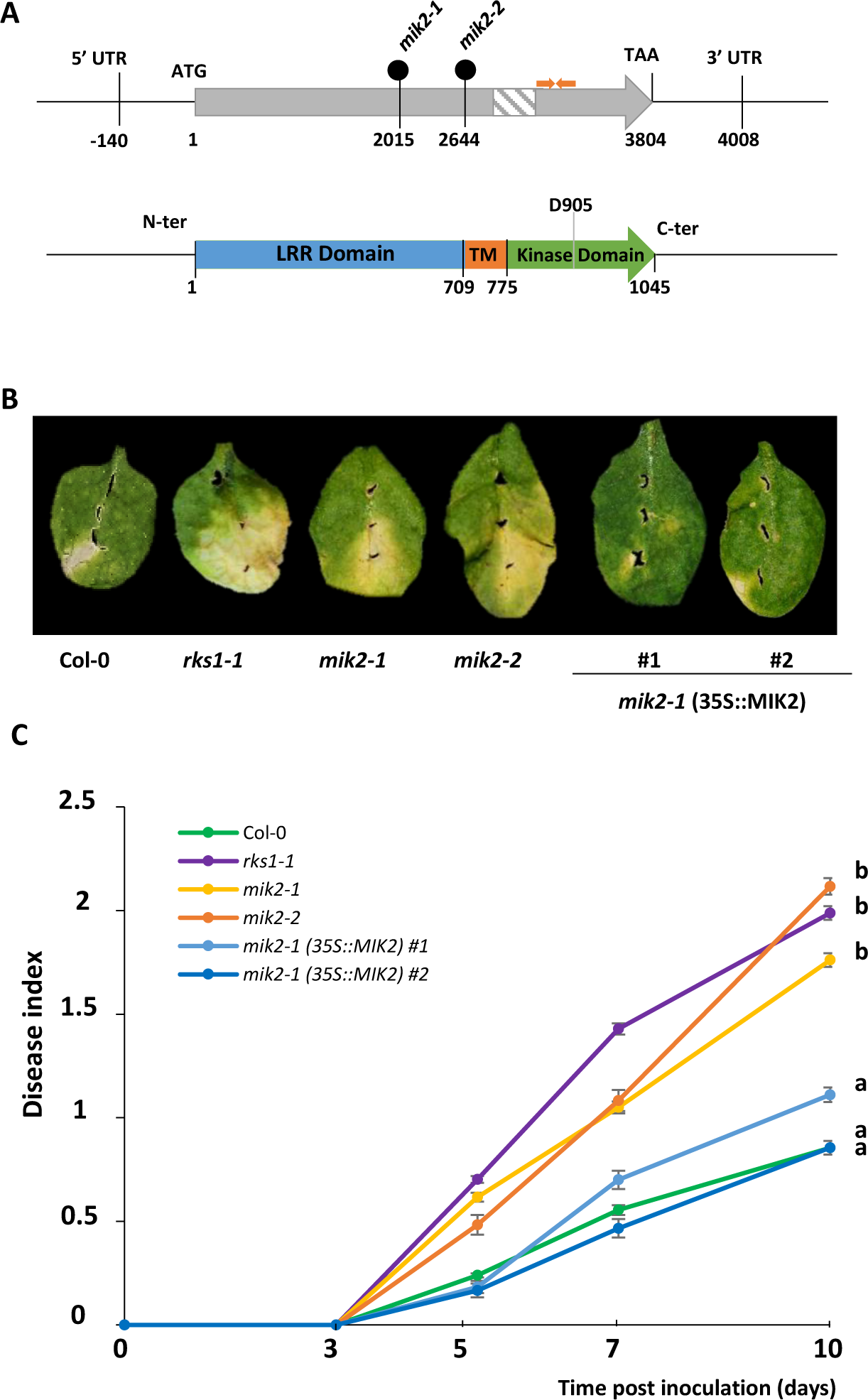
*mik2* mutants are more susceptible to *Xcc568* inoculation. **A,** Schematic representation of the genomic organization of the *mik2* mutants and of the MIK2 protein. *MIK2* consists in two exons (grey boxes) and one intron (hatched box). For *mik2-1* and *mik2-2*, the T-DNA is integrated in the first exon, respectively at the position 2015 and 2644. The orange arrows indicate the primers used for measurement of *MIK2* gene expression. For the schematic representation of the MIK2 protein, blue represents the Leucine-Rich Repeat (LRR) domain; orange, the transmembrane domain (TM); green, the kinase domain and D905 the aspartic amino acid involved in the kinase catalytic site. **B,** Disease symptoms were observed on leaves of *mik2* and *rks1* mutants, two *mik2-1* complemented lines with a *35S::MIK2* construct (#1 and #2) and Col-0 wild-type plants, 10 days post-inoculation with a bacterial suspension adjusted to 2.10^8^ cfu/mL. **C,** Time course evaluation of disease index after inoculation of *rks1-1* (purple), *mik2-1* (yellow), *mik2-2* (orange), *mik2-1 (35S::MIK2) #1* (light blue), *mik2-1 (35S::MIK2) #2* (dark blue) and the wild type Col-0 (green) with *Xcc568* under the same conditions as B. Means and standard errors were calculated from 5 plants and based on 3 independent experiments. Different letters indicate significant differences among lines for disease index kinetics.

To further explore the respective roles of *MIK2* and *RKS1* in the QDR, we generated the double mutant *rks1-1/mik2-1* and phenotyped it in response to inoculation with *Xcc568* (Fig. 3A and 3B, Supplemental Fig. S3). The double mutant was significantly more susceptible than Col-0, slightly more susceptible than the single mutants, and less susceptible than the susceptible reference accession Kas-1. This result suggests that *MIK2* might at least be partially involved in the *RKS1* pathway mediating QDR to *Xcc568*. To investigate more precisely the respective roles of *MIK2* and *RKS1* in these pathways, we analyzed the gene expression of a total of 36 genes highly connected to RKS1 from the previously reconstructed *RKS1*-dependent gene network (Delplace *et al*., 2020) or described to be dependent from *MIK2* (Hou *et al*., 2021). Expression of these genes was measured by RT-qPCR in Col-0 and the mutants *rks1-1*, *mik2-1*, *mik2-2*, *rks1-1/mik2-1*, 6 hours after inoculation by *Xcc568* (Supplemental Table 1). A significant fraction of genes (44%) showed an expression dependent of the mutation *mik2* (either *mik2-2,* or both *mik2-1* and *mik2-2)*. In this fraction, 31% showed an expression profile dependent of both mutations *rks1* and *mik2*. Several examples are presented in Fig. 3C. These results clearly support the hypothesis that *MIK2* and *RKS1* initiate common signaling pathways to control gene expression. However, these common pathways appear to constitute only a fraction of the ones initiated by both proteins, in good agreement with the double mutant phenotype reported in Fig. 3A and B. Interestingly, *MIK2* gene expression was significantly reduced in two independent *RKS1* overexpressing lines (*RKS1-OE1* and *RKS1-OE2*) (Huard-Chauveau *et al*., 2013; Delplace *et al*., 2020) (Supplemental Fig. S3B), suggesting that *MIK2* gene expression might be regulated by an overexpression of *RKS1*.

**Figure 3.**
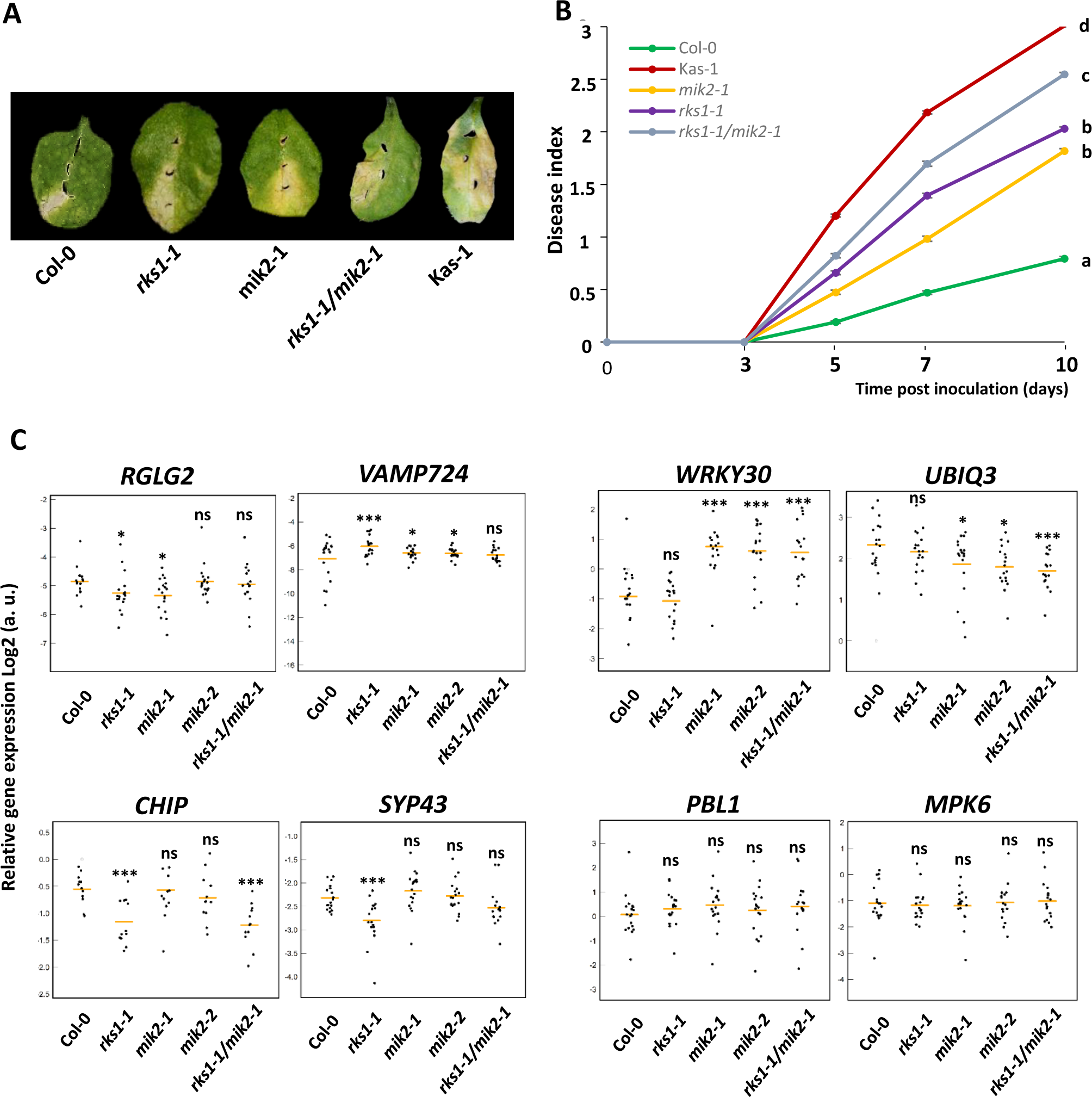
*MIK2* participates to the *RKS1* dependent control of QDR in response to *Xcc*. **A,** Disease symptoms were observed on leaves of *mik2-1* and *rks1-1* mutants, the double mutant *rks1-1/mik2-1*, the susceptible accession Kas-1 and the wild-type Col-0, 10 days after inoculation with a bacterial suspension of *Xcc568* adjusted to 2.10^8^ cfu/mL. **B,** Time course evaluation of disease symptoms and index after inoculation of *mik2-1* (yellow), *rks1-1* (purple), *rks1-1/mik2-1* (grey), the susceptible accession Kas-1 (red) and the wild type Col-0 (green) with a bacterial suspension of *Xcc568* adjusted to 2.10^8^ cfu/mL. Disease scores were observed at 3-, 5-, 7- and 10-days post-inoculation. Means and standard errors were calculated from five plants based on six independent experiments. Different letters indicate significant differences among lines for disease index kinetics. **C,** Jitter plots illustrating RT-qPCR results of the expression profile of genes from the RKS1-dependent gene network (Delplace *et al*, 2020), 6 hours after inoculation with *Xcc568* (2.10^8^ cfu/mL) in leaves of the wild type accession, Col-0, in *mik2-1* and *mik2-2* mutants, the *rks1-1* mutant and *rks1-1/mik2-1* double mutant. Dots correspond to relative gene expression value adjusted for differences between the two experiments. Mean relative gene expression is represented by an orange segment. The *RGLG2* and *VAMP724* genes present an expression profile dependent of the presence of *MIK2* and *RKS1*. The *WRKY30* and *UBIQ3* genes present an expression profile dependent of the presence of *MIK2* and independent of *RKS1*. The *CHIP* and *SYP43* genes present an expression profile independent of the presence of *MIK2* and dependent of *RKS1*. The *PBL1* and *MPK6* genes present an expression profile independent of the presence of *MIK2* and *RKS1*. Statistical results compared to Col-0 were generated with a GLM procedure in the *R* environment and based on three independent experiments with six plants/line/experiment. ns: non-significant, * *P* < 0.05, ** *P* < 0.01, *** *P* < 0.001. A correction for the number of tests was performed to control the False Discovery Rate (FDR) at a nominal level of 5%.

### Catalytic activity of MIK2 is required for QDR to *Xcc568*

Taking in account that (i) MIK2 participates to the *RKS1* dependent QDR in response to *Xcc*, and (ii) RKS1 physically interacts with the kinase domain of MIK2, we tested whether these MIK2 functions depend on its kinase catalytic activity. For this purpose, an inactive catalytic mutant version of MIK2 full-length protein, MIK2^D905A^, affected in the putative active HRD site of the MIK2 kinase domain, was generated. Kinase assays using MIK2 and MIK2^D905A^ showed that MIK2 was able to autophosphorylate and that the aspartic acid residue in position 905 was required for MIK2 autophosphorylation activity (Fig. 4A). Firstly, the catalytic mutated version of the kinase domain of MIK2, MIK2-KD^D905A^-HA, was able to interact with RKS1^D191A^ fused to the c-myc tag, as shown for the wild type version of MIK2 (Fig. 1). After immunoprecipitation with an anti-HA antibody, RKS1^D191A^-c-myc was detected when incubated with MIK2-KD^D905A^-HA (Fig. 1A), indicating that the acid aspartic residue of the HRD motif of MIK2 is not required for the RKS1-MIK2 interaction. Furthermore, incubation of MIK2 and MIK2^D905A^ protein with RKS1 or RKS1^D191A^ did not reveal MIK2 or RKS1 phosphorylation activities *in vitro* (Fig. 4A). Secondly, after introduction of MIK2^D905A^ into the *mik2-1* mutant for a complementation test, three transgenic lines were characterized and inoculated with *Xcc568* (Supplemental Fig. S3C). The three complemented lines showed significant phenotypic differences from Col-0, but similar phenotypes to the *mik2-1* mutant (Fig. 4B and 4C). MIK2^D905A^ version was therefore not able to restore resistance in the *mik2-1* mutant at a similar level as the wild type. These results indicate that the catalytic activity of MIK2 is required to confer QDR to *Xcc568.* Finally, transgenic lines overexpressing *MIK2* or *MIK2^D905A^* in Col-0 were also characterized (Fig. 4D). *MIK2* expression was increased in 35S::MIK2 lines (517% and 161% in the two lines compared to Col-0) and reduced in 35S::MIK2^D905A^ lines (26% and 42% in the two lines compared to Col-0) (Supplemental Figure S3E). Inoculation tests of these lines revealed that MIK2 and *MIK2^D905A^* mis-expression in Col-0 did not significantly affect resistance to *Xcc568*, with the exception of one line in the case of *MIK2* overexpression for which a slight reduction of resistance to *Xcc568* was observed (Fig. 4D).

**Figure 4.**
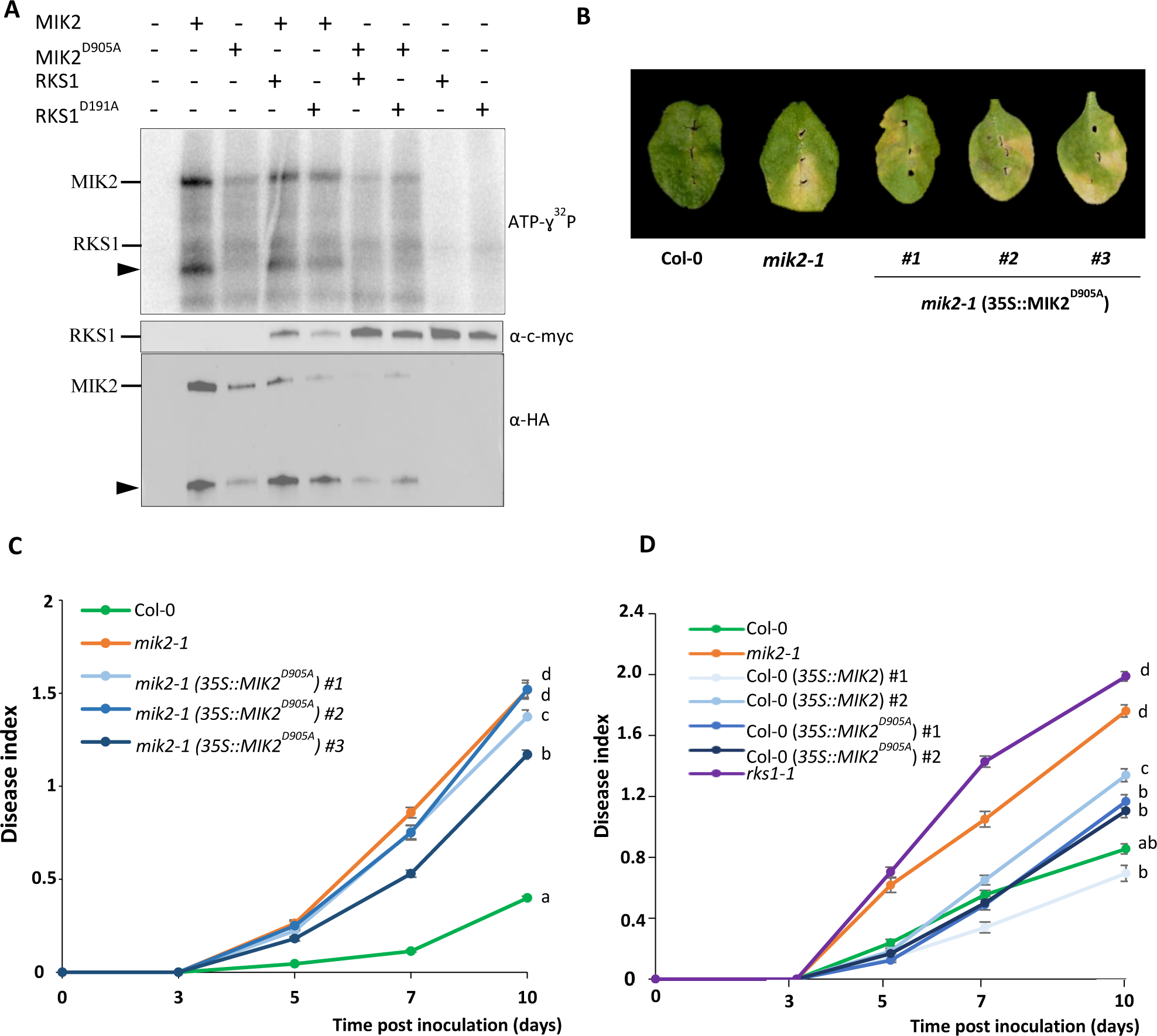
The catalytic activity of MIK2 is required for resistance to *Xcc568* but overexpression of MIK2 and MIK2*^D905^* in Col-0 does not increase resistance to *Xcc568*. **A,** Kinase assay with protein extracts from *N. benthamiana* leaves transformed by Agrobacterium carrying the constructs MIK2-HA, MIK2^D905A^-HA, RKS1-c-myc or RKS1^D191A^-c-myc. The total protein extracts were subjected to anti-HA or anti-c-myc immunoprecipitation. The immunoprecipited proteins were incubated for 2 hours at room temperature with ATP-ɣ^32^P. The proteins were separated on a 4-15% SDS-PAGE gel and detected by anti-HA, anti-c-myc immunoblot and the presence of ATP-ɣ^32^P by autoradiography. Black triangles indicate the potential MIK2 kinase domain cleaved. **B,** Disease symptoms were observed on leaves of *mik2-1* mutant, three *mik2-1* complemented lines with a 35S::MIK2^D905A^ construct and wild-type plants, 10 days post-inoculation with a bacterial suspension of *Xcc568* adjusted to 2.10^8^ cfu/mL. **C,** Time course evaluation of the disease index of *mik2-1* mutant (orange), three *mik2-1* complemented lines with a 35S::MIK2^D905A^ construct (blue) and wild-type plants (green) after inoculation with *Xcc568* under the same conditions as A. Means and standard errors were calculated from 5 plants per line per experiment and based on 3 or 4 independent experiments. a, b, c or d represent statistical groups based on comparison of disease index kinetics. **D,** Time course evaluation of the disease index of lines overexpressing *MIK2* or *MIK2^D905A^* in Col-0 (35S construction), *rks1-1* and *mik2-1* mutants and the wild type plants inoculated with a bacterial suspension of *Xcc568* adjusted to 2.10^8^ cfu/mL. Means and standard errors were calculated from 5 plants per line and based on 3 independent experiments. a, b, c or d represent statistical groups based on comparison of disease index kinetics.

### MIK2 and RKS1 take place in an RLK network

We demonstrated that MIK2 and RKS1 form a complex that is involved in QDR to *Xcc*. It is now tempting to speculate, in view of (i) our results showing that these two proteins share only a part of their downstream signaling pathways, (ii) the RKS1 dependent network previously identified (Delplace *et al*., 2020), and (iii) the literature data describing MIK2 as a multiple ligands perceiving receptor, that these two proteins do not act isolated and require other perception/signalling proteins to orchestrate QDR. To test this hypothesis, we first identified from the literature, proteins experimentally demonstrated to physically interact with MIK2. A MIK2-centralized network was reconstructed including 25 RLKs, BSK3 and RKS1. Nine and five RLKs interacting with MIK2 were described in literature to be involved respectively in plant immunity (Kemmerling *et al*., 2007; Blaum *et al*., 2014; Mata-Pérez *et al*., 2015; Liu *et al*., 2016; Yeh *et al*., 2016; Mendy *et al*., 2017; Burgh *et al*., 2019; Chan *et al*., 2020; Laohavisit *et al*., 2020) and plant development (Takeuchi and Higashiyama, 2016; Wang *et al*., 2016; Duckney *et al*., 2017; Crook *et al*., 2020). Interestingly, seven proteins from the RKS1 dependent protein – protein network were found highly connected to the MIK2-centralized network (Fig. 5A). Given that (i) proteins with an extracellular domain (ECD) length up to 400 amino acids may act as putative co-receptors, and (ii) proteins with a larger ECD may act as putative ligand recognition receptors (Xi *et al*., 2019), this network is composed of 25 RLKs described as ligand perception proteins (including MIK2) and 12 co-receptors possibly implicated in signal transduction events (Fig. 5B).

**Figure 5.**
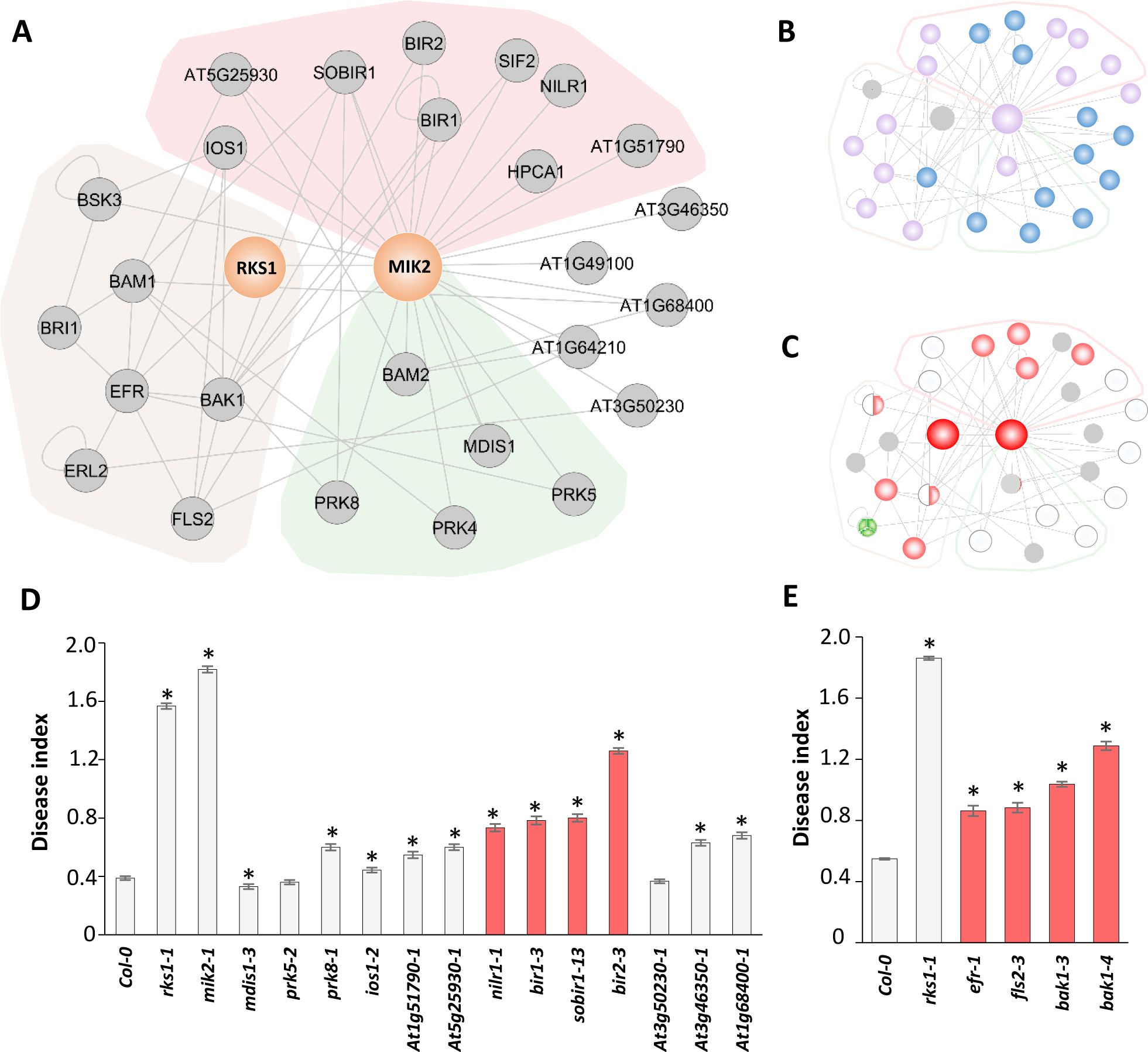
MIK2 and RKS1 as hubs of an RLK network. **A,** MIK2 protein–protein interaction subnetwork was reconstructed from STRING and BioGrid databases and plotted with Cytoscape. The light brown area indicates proteins recovered from the RKS1 dependent network (Delplace *et al*., 2020). The red area indicates RLKs interacting with MIK2 and described to play a role in plant immunity (Kemmerling *et al*., 2007; Blaum *et al*., 2014; Mata-Pérez *et al*., 2015; Liu *et al*., 2016; Yeh *et al*., 2016; Mendy *et al*., 2017; Burgh *et al*., 2019; Chan *et al*., 2020; Laohavisit *et al*., 2020). The green area indicates RLKs interacting with MIK2 and described to be implicated in plant development (Takeuchi and Higashiyama, 2016; Wang *et al*., 2016; Duckney *et al*., 2017; Crook *et al*., 2020), the uncolored zone corresponds to hypothetical RLKs. **B,** In the MIK2 protein-protein interaction network, purple circles represent RLKs from the “ligand perceiving group” (Xi *et al*., 2019), blue circles represent LRR-RKs from the “co-receptor group” and grey circles, other proteins.**C,** Each circle represents the phenotype of the mutant corresponding to the protein component of the network. Colored circles indicate some mutants significantly affected in their response to *Xcc568*. Mutants represented in red are significantly affected in their response to *Xcc* (at least 30% of increased susceptibility compared to Col-0 disease index), according to Fig. 5D and 6E, and those represented in green are more resistant compared to Col-0 according to Delplace *et al*. (2020). The circle is divided according to the number of tested mutants. White circles correspond to mutants with a phenotype similar to the one of the wild type line. No mutant was tested for genes represented in grey. Data presented for ERL2 and BSK3 were recovered from Delplace *et al*. (2020). **D,** Disease index of mutants corresponding to genes belonging the MIK2-RKS1 network, at 10 dpi after inoculation with a bacterial suspension adjusted to 2.10 cfu/mL. *Represents kinetic modeling difference with Col-0 time course, based on five independent experiments with 5 plants / mutant. Light grey bars correspond to mutants for RLKs described in part A and dark grey bars correspond to *rks1-1* mutant and Col-0 wild type used as controls. Mutants represented in red are significantly affected in their response to *Xcc* (at least 30% of increased susceptibility compared to Col-0 disease index). **E,** Disease index at 10 dpi after inoculation in the same conditions as D.*represents kinetic modeling deference with Col-0 time course in three to five experiments with 5 plants / mutant. Light grey bars indicate mutants for RLKs from the RKS1 dependent network and dark grey bars correspond to *rks1-1* mutant and Col-0 wild type used as controls. Mutants represented in red are significantly affected in their response to *Xcc* (at least 30% of increased susceptibility compared to Col-0 disease index).

In order to evaluate the possible implication of some of the MIK2-RKS1 network components in QDR to *Xcc*, 17 mutants (corresponding to 16 genes) were collected and characterized for their response to *Xcc568*. The phenotypic data are summarized in Fig. 5C to E. Eight mutants corresponding to seven receptors showed a strong and significant alteration in their response to *Xcc568* (at least 30% of increased susceptibility), compared to Col-0, while mutants corresponding to seven other receptors showed a weak but significant alteration (Fig. 5D and 5E) (Supplemental Table 2 and 3). Interestingly, mutants showing a strong alteration correspond to network components known to operate in plant immunity, while the mutants with a weak effect correspond to components associated with plant development or without known function. Because MIK2 was recently proposed to operate by sensing cell wall perturbations (Van der Does *et al*., 2017), we also phenotyped several cell wall integrity altered mutants, *i.e. herk1*, *the1-4*and *FER* (Cheung and Wu, 2011; Nissen *et al*., 2016), in response to inoculation with *Xcc* (Fig. 3). *herk1-1* and *the1-4* mutants did not show a significantly altered response to *Xcc568,* while *fer1-5* mutants were slightly more resistant than Col-0. Our data suggest that the genes *HERK* and *THE1* are not significantly involved in QDR to *Xcc*. In addition, mutants for *MIK1* and *MIK2-like*, the closest homologs of *MIK2*, show only weak alteration in response to *Xcc* (Fig. 3). Taken together, these results indicate that at least seven RLKs, highly connected to MIK2 and/or RKS1, have a partial effect in QDR to *Xcc568.* These findings reveal the complexity of the molecular mechanisms involved in MIK2/RKS1-dependent signalling pathways, in good agreement with the genetic and molecular complexity of QDR.

## Discussion

Even if QDR is the most prevalent form of resistance in natural populations and crop fields, the identification of genes involved in the molecular mechanisms underlying QDR is still in its early stages (Roux *et al*., 2014b; French *et al*., 2016). In this study, we have identified the LRR receptor kinase MIK2 as an interactor of RKS1, a major determinant of QDR to the bacterial pathogen *Xcc* (Huard-Chauveau *et al*., 2013; Delplace *et al*., 2020). In addition, we found that MIK2 plays a role in QDR to *Xcc*, depending on its kinase catalytic activity. Taking in account (ii) the RKS1 dependent network previously identified (Delplace *et al*., 2020), and (ii) the ability of MIK2 to interact with diverse RLKs (receptors and co-receptors, Xi *et al*., 2019), themselves involved in interaction with RKS1 protein partners, our results suggest a role for the complex MIK2-RKS1 as a scaffolding hub in a RLK network to modulate Xanthomonas determinant perception for QDR signaling and activation. The functional relevance of this RLK network, centered on the MIK2-RKS1 complex, has been addressed by mutational analysis, thereby revealing the involvement of several RLK components in QDR regulation. While the molecular mechanisms underlying QDR remain poorly characterized, combining functional validation, protein-protein interaction analysis and network reconstruction, revealed new insights into a perception system that regulates QDR in response to *Xcc*.

### The cell surface receptor-like kinase MIK2 physically interacts with RKS1 to activate QDR

By using the yeast two hybrid system, we previously identified 43 proteins as putative interactors of RKS1, including 21 metabolism related proteins and 6 signaling related components (ex: EFR, KIN11, SGT1a, MKP1, RANBP1 and NLP7) (Delplace *et al*., 2020). Here, we demonstrated the interaction between RKS1 and the kinase domain of the receptor-like kinase MIK2. Co-localization and BiFC experiments confirmed the subcellular localization of the two proteins and their interaction in the plasma membrane. Interestingly, RKS1 is localized in the plasma membrane, cytoplasmic tracks and the nucleus (Delplace *et al*., 2020), suggesting different functions for RKS1 including a contribution to pathogen perception events. The NLR ZAR1 was demonstrated to form a complex with RKS1 in the cytoplasm to perceive the injected effector AvrAC from *X. campestris* (Wang *et al*., 2019; Adachi *et al*., 2019). The interaction of RKS1 with a receptor like protein such as MIK2, suggests that RKS1 might also be involved in the extracellular perception of pathogen determinants at the plasma membrane. Based on our findings, RKS1 might be part of membrane complexes that, similarly to MIK2, include cell surface receptors required for transduction events (Antolín-Llovera *et al*., 2012). Similar to the pseudokinase domain of the receptor BIR2 involved in the allosteric regulation of the receptor BAK1 (Blaum *et al*., 2014), the RKS1-MIK2 complex, or RKS1, could act as a scaffolding protein complex and participate to the regulation of signaling pathways (Roux *et al*., 2014a; Murphy *et al*., 2017). Interestingly, MIK2 seems connected with multiple sensing systems including cell wall integrity sensing (CWIS) (Rhodes *et al*., 2021), root growth and response to abiotic and biotic stresses (Julkowska *et al*. 2016; Van der Does *et al*., 2017). In particular, MIK2 is involved in the perception of (i) the peptide LURE1, a chemotactic attractant of the pollen tube, by heterodimerizing with MDIS1 and MDIS1-Interacting Receptor-like Kinase 1 (Wang *et al*., 2016) and (ii) the phytocytokine peptide SCOOP12, by heterodimerizing with BAK1 (Rhodes *et al*., 2021). While no plant developmental function has been suggested for RKS1 yet, it is a major determinant of QDR for which cell wall integrity sensing might be components (Zuo *et al*., 2015b; Bani *et al*., 2018). To test this hypothesis, mutants for CWIS (*herk1-1*, *the1-4* and *fer1-5*) were phenotyped in response to *Xcc*. None of the CWIS mutants was found more susceptible to *Xcc568* (Fig. S4), indicating that CWIS is not required to confer QDR to *Xcc*. These results show that CWIS does not generally interfere with QDR to *Xcc*, in opposition with a previous study proposing that *MIK2* and *THE1* could function in the same pathways but distinct from the *FER* pathways, against *Fusarium oxysporum* (Coleman *et al*., 2020). This discrepancy might be explained by the fact that *F. oxysporum* is more a root and cell wall invasive pathogen in comparison with the leaf vascular pathogen *Xcc* (Kubicek *et al*., 2014; An *et al*., 2019). MIK2 participates to detection of multiple ligands (Wang *et al.,* 2016, Rhodes *et al.,* 2021), that could require directly MIK2 or MIK2 interacting with other ligand-perceiving receptors (Fig. 5B). In the future, to decipher the molecular dialogues between *Xcc* and host plants, a challenge would be to identified putative *Xcc* or/and plant cell damages derived ligands perceived by MIK2. Approaches including physical protein-ligand interactions using MIK2 and other potential RLKs identified in this study, would help to shed some light on the perception events responsible for QDR. To conclude, *MIK2* with its diverse functions in CWIS and responses to abiotic and biotic stress, could play different functional roles depending of its interacting partners. As a complex with RKS1, it appears to be a major player in plant immune responses to the bacterial pathogen *Xcc*.

### RKS1-MIK2 as a regulatory complex for QDR signaling?

While receptor kinases, and among them the LRR-receptor kinases, constitute the largest protein kinase family in plants, a large part of these receptors and their putative interactors are still functionally uncharacterized. Here, we showed that *MIK2* confers QDR to *Xcc568* at a similar level as *RKS1*. Analysis of the double mutant *rks1-1/mik2-1* revealed that *MIK2* and *RKS1* probably act in a common pathway in response to *Xcc,* which might represent only a part of the immune response, as the double mutant express an intermediary phenotype, more susceptible that the wild type resistant accession, less susceptible than the susceptible reference accession, but slightly more susceptible than the single mutants. To further explore their respective functions, we measured the expression of a set of 36 genes from the previously identified RKS1 dependent network (Delplace *et al*., 2020) or described to be dependent from *MIK2* (Hou *et al*., 2021). A major part of these genes (44%) showed an expression dependent of the mutation *mik2*. For this set of genes, 31 % (including *RGLG2* and *VAMP724,* presented in Fig. 3) showed an expression profile dependent of both mutations *rks1* and *mik2*, confirming the existence of common pathways activated by both proteins. *RGLG2* was described as a RING domain ubiquitin E3 ligase implicated in stress drought adaptation and VAMP724 as SNAREs protein regulated vesicle-associated trafficking (Yu *et al*., 2021; He *et al*., 2022). Eleven genes were found specifically dependent of *MIK2*, such as the transcription factor WRKY30, implicated in abiotic stress tolerance (El-Esawi *et al*., 2019). These different roles can be explained by the putative interactions of MIK2 with other signaling partners in plasma membrane receptor complexes (He *et al*., 2018). Together these data confirm the implication of *MIK2* and *RKS1* in QDR signaling, potentially *via* the MIK2 kinase domain. Indeed, we showed that MIK2, as classical RD (Arginine, Aspartic acid) RLKs like BAK1 and CERK1 (Oh *et al*., 2010; Suzuki *et al*., 2018), exhibits an auto-phosphorylation activity dependent on aspartic acid 905 residue. Similar to BRI1 that is maintained in an inactive stage by auto-phosphorylation (Oh *et al*., 2012), MIK2 phosphorylation activity could be required for auto-regulation. Surprisingly, the MIK2 aspartic acid 905 residue appeared not essential for RKS1 interaction. These results suggest, in agreement with the functioning of multiple plasma membrane receptor complexes (Burkart and Stahl, 2017), that RKS1 might act as a scaffolder for receptor complexes assembly in the context of QDR. A similar role was described for the PTI receptors BAK1, FLS2 and EFR, with the FER scaffolding receptor (Stegmann *et al*., 2017). Our results are in good agreement with the recent “Invasion model” proposing that there might be no clear distinction between PTI, ETI and more widely the different forms of plant immunity responses (Cook *et al*., 2015; Kanyuka and Rudd, 2019) (including QDR), and that broad-spectrum immunity might depend on complex networks of cell surface immune receptors in most cases. Here, we extend this model by proposing that RKS1 participates to surface and intracellular perception mechanisms (in interaction either with MIK2 or ZAR1), which are initiated either in the apoplast or the cytosol.

### MIK2/RKS1 complex as a hub to coordinate a RLK perception network?

LRR-RLKs sense a wide array of molecules exogenously or endogenously produced, and regulate plant growth and immunity. They can operate in a regulatory network (Smakowska-Luzan *et al*., 2018) where small LRR-RKs (co-receptors) are involved (i) in fine-tuning (by activation or stabilization) of ligand binding receptors, and (ii) as regulatory scaffolds for the organization of the signaling network (Xi *et al*., 2019). In this context, understanding the functioning of the MIK2/RKS1 complex in QDR needs to be replaced within a protein–protein interaction (PPI) network. We identified from the literature MIK2 interacting proteins, and generate a network including beyond RKS1, RLKs from the RKS1 dependent network. MIK2, a ligand binding receptor, takes place in a highly connected RLK network including signaling proteins involved either in immunity or plant development (Fig. 5). Interestingly, the MIK2-RKS1 network reveals no clear co-receptor hub (linked to ligand-perceiving receptors) such as BAK1 (Smakowska-Luzan *et al*., 2018). In addition, MIK2 is connected to diverse receptors and not only with co-receptors, such as the ligand-perceiving receptor BAM1 that forms a complex with several co-receptors of the CIK-family (Cui *et al*., 2018). The MIK2-RKS1 centralized network appears more diverse than co-receptor and ligand perceiving receptor associations (Xi *et al*., 2019). This might be due to the dynamic complexity of the perception events involved in QDR, and probably also to the insufficient characterization / absence of some LRR-RLKs (or other receptors) in this reconstructed network. A functional analysis of this network by mutant analysis showed that seven RLKs are involved in QDR to *Xcc* with a strong effect (Fig. 5), described to be related to plant immunity and connected with MIK2 (BIR1, BIR2 NIRL1, SOBIR, EFR, FLS2 and BAK1) (Kemmerling *et al*., 2007; Blaum *et al*., 2014; Liu *et al*., 2016; Laohavisit *et al*., 2020; Rhodes *et al*., 2021). Among these RLKs, the MIK2/RKS1 complex might recruit some RLKs as a perception system to trigger QDR, similarly to the co-receptor BAK1. RKS1 can also be required for transduction events after MIK2-RLK complex formation. Seven other RLKs connected with MIK2, including co-receptors with unknown functions (*AT1G68400* and *AT1G64210*) or with plant developmental functions (MDIS1 and PERK8) are implicated in QDR to *Xcc* but to a much lesser extent, indicating that MIK2 in association with RKS1, could mainly recruit RLKs involved in immune responses. Similar to the receptor BAK1 described to be a molecular switch between PTI and plant development (Postel *et al*., 2010; Zhou *et al*., 2019), MIK2 might require specific RLKs or other signaling components to coordinate immunity and development perception systems. Interestingly, cytoplasmic kinases are also known as molecular switches between plant development and immunity pathways, such as BSKs (brassinosteroid kinases) (Lin *et al*., 2013). BSK3, which is implicated in brassinosteroid signaling, confers partial resistance to *Xcc* and thus can be a key regulatory compound in the MIK2/RKS1-centralized network (Sreeramulu *et al*., 2013; Delplace *et al*., 2020). In this line, BSK3 was also described as a scaffolding protein for plant development (Ren *et al*., 2019). Another interesting feature of this network is the presence of four RLKs from the RKS1 dependent network: ERL2, BAK1, EFR and FLS2. EFR and FLS2 are known to directly perceive pathogen determinants (Chinchilla *et al*., 2006; Zipfel *et al*., 2006) and can therefore participate to the perception of multiple pathogen determinants, which is in good agreement with the view of QDR as a complex network integrating diverse pathways in response to multiple pathogenic determinants (Roux *et al.,* 2014b). While receptors could participate directly or indirectly to pathogen perception with strong implication in QDR to *Xcc*, like BAK1 or BIR2, other receptors with weak implication in QDR, like MDSI1 or IOS1, could have either redundant functions with other receptors, or only an indirect implication in *Xcc* perception. In the future, consolidation of the PPI network, investigation of other possible interactions among the different components and test if the contribution of these RLKs in resistance to *Xcc* depends on the presence of RKS1 or MIK2, will permit to confirm and extend the complexity of this network. Taken together, our results suggest that the RKS1/MIK2 complex in relation with a number of additional RLKs might participate to a perception system and act as a scaffolder hub to coordinate assembly of multiple perception complexes.

In summary, the newly demonstrated MIK2/RKS1 complex takes place in a RLK network in the plasma membrane and might be involved in perception of multiple pathogen ligands, subsequently leading to QDR, each component participating partially to QDR to *Xcc*. This work constitutes a strong basis for deciphering molecular mechanisms involved in the perception systems related to QDR.

## Material and methods

### Plant material

Arabidopsis plants were grown on Jiffy pots under controlled conditions (Lacomme and Roby, 1996), in a growth chamber at 22 °C with a 9-h photoperiod at 192 µmol m^-2^ s^-1^. We used the wild-type line Columbia (Col-0) (Wilson *et al*., 2001; Li *et al*., 2006) and the following mutant lines (all in Col-0 background): *rks1-1* (Huard-Chauveau *et al*., 2013), other mutants from the GABI-kat (http://www.gabi-kat.de) or SALK (http://signal.salk.edu) seed libraries and the susceptible accession to *Xcc568* Kashmir-1 (Kas-1) (Huard-Chauveau *et al*., 2013). For each mutant line, the T-DNA insertion was determined by sequencing of the T-DNA borders (GABi o8409 or LBb1.3_SALK primer) and of the flanking regions (Supplemental Table 4). The double mutant *mik2-1/rks1-1* was obtained by crossing *mik2-1* pistils with *rks1-1* pollen. For transient expression assays, *N. benthamiana* plants were cultivated 4-week at 21°C and under 15 hour light period/ 9 hour dark period.

### Bacterial material

*Agrobacterium tumefaciens* strains GV3101 and C58C1 were grown at 28°C on YEB medium with 50 µg.mL^-1^ rifampicin, 10 µg.mL^-1^ kanamycin, complemented with 10 µg.mL^-1^ gentamicin (GV3101) or tetracyclin (C58C1). For transient expression assays for fluorescence microscopy, the Agrobacterium strain GV3101 carrying the corresponding mRFP1 and eGFP constructs was used. Arabidopsis seedlings were transformed according to Marion *et al*. (2008). Overnight cultures of *Agrobacterium tumefaciens* were resuspended in 5% sucrose supplemented with acetosyringone (200 µM). Then one week-old seedlings were vacuum-infiltrated with the *Agrobacterium* solution. For co-immunoprecipitation experiments,

*Agrobacterium* strain C58C1 carrying the corresponding HA and c-myc constructs was prepared as GV3101 strain. *N. benthamiana* plants were infiltrated and leaf disks harvested 28h after inoculation.

*Xcc* inoculation tests were done with the strain LMG568/ATCC33913 (*Xcc568*) (Da Silva *et al*., 2002). Cultures of *Xcc568* were grown at 28°C on Kado medium (Kado and Heskett, 1970) supplemented with 50 µg.mL^-1^ rifampicin and 25 µg.mL^-1^ kanamycin.

### Constructs and plant transformation

The *MIK2* full length constructs were performed by amplification of the *MIK2* (*AT4G08850*) cDNA coding sequence using respectively [attb1_MIK2_full_length and attb4_MIK2_NO_STOP] as primers (Supplemental Table 5). The catalytic MIK2^D905A^ mutants were generated using respectively [MIK2_D905A_fw and MIK2_D905A_rev] primers (Supplemental Table 5). MIK2-KinaseDomain was amplified with [attb1_MIK2_KD_fw and attb4_MIK2_NO_STOP] primers (Supplemental Table 5). PCR products were cloned into the multisite Gateway entry vector pBS-DONR P1-P4. 3xHA, 5xc-myc, mRFP1 and eGFP tags were cloned into the entry vector pBS-DONR P4-P2. To fuse *MIK2* constructs with 3xHA, 5xc-myc, mRFP1 or eGFP tags, both vectors were mixed with the 35Sp plant expression vector pEarleyGate100 (Lema Asqui *et al*., 2018) and recombined with LR clonase II (Invitrogen) described previously (Gu and Innes, 2011). We used RKS1^D191A^ in pBS-DONR P1-P4 described in Delplace *et al*. (2020) fused as MIK2 with c-myc in pEG100. The plasmid constructs with the different tags fused to the genes of interest are reported in the Supplemental Table 6.

The *MIK2-OE/Col-0* and *MIK2-OE/mik2-1* constructs were introduced in the C58C1 *Agrobacterium* strain for transformation of Arabidopsis Col-0 (Clough and Bent, 1998). Selection of transformed plants were performed by spraying glufosinate ammonium (BASTA) at 10mg.L^-1^ on soil-grown plants. Harvested seeds were spread on MS medium containing 50 μM of phosphinothricin for selection of homozygous transgenic plants.

### Plant phenotyping and statistical analyses

The disease symptoms of mutant lines were evaluated after inoculation with a bacterial suspension adjusted to 2.10^8^ cfu.mL^-1^, in three independent experiments (Lacomme and Roby, 1996), as compared to Col-0 and *rks1-1*. Four leaves per plant and four 28-day old plants per line were inoculated by piercing and scored as previously described (Meyer *et al*., 2005), at 0, 3, 5, 7 and 10 days post inoculation.

We fitted the temporal relation of disease index between the tested mutant line and Col-0 by a second order polynomial using the *lm* library under the *R* environment (https://www.R-project.org/). Kinetics of disease index were considered similar if the coefficient of the second order was not significantly different from 0 and if the slope was not significantly different from *P*-value numbers represent kinetic modeling deference with Col-0, 0 = *P* > 0.05 and 1= *P* ≤ 0.05.

### RNA isolation and q-RT-PCR

RNA extraction was performed with NucleoSpin***®*** RNA plus kit (Macherey-Nagel). RNA extraction was performed with leaves from 28-days-old healthy plants (6 plants, 3 leaves/plant) or inoculated with the *Xcc568* (6h post inoculation) or treated with the XOOP14 peptide at 1µM (3h post-treatment). RT-qPCR analysis was performed as described by Khafif *et al,*. (2017). The gene *AT2G28390* (*MONENSIN SENSITIVITY 1)* was used as an internal control as it is known to be stable in our physiological conditions (Czechowski *et al*., 2005). Average ΔCp was calculated from three experiments and data were expressed as fold induction for each point as compared to the wild type. Primers have been designed *via* the Roche website (http://www.lifescience.roche.com) (Supplemental Table 7 and 8). Results were analyzed using the LC480 on-board software, release version 1.5.0.39. Statistical results were generated with a GLM procedure under the *R* environment. A correction for the number of tests was performed to control the False Discovery Rate (FDR) at a nominal level of 5%.

### Co-immunoprecipitation assays

Total proteins were extracted from *N. benthaniama* leaves harvested 28h after treatment (transient expression experiments) using the extraction buffer [10% glycerol, 50mM Tris-HCl pH 8, 150mM NaCl, 1mM DTT, 0.05% Nonidet P-40, cOmplete Mini EDTA-Free protease inhibitor cocktail (Sigma), PhosSTOP EASYpack phosphatase inhibitors (Roche) and a spatula tip of polyvinylpolypyrroidone (PVPP)]. The samples were incubated with anti-c-myc or anti-HA magnetic beads (Thermo Scientific) for 2-4 hours and washed with the same buffer without phosphatase inhibitors. Proteins were then separated on SDS-PAGE 4-15%. After incubation with rat anti-HA:HRP (3F10 clone, Roche [dilution 1:3000]) or mouse anti-Myc:HRP antibodies (9E10 clone, Roche [dilution 1:3000]), proteins were visualized using the Bio-Rad Clarity™ Western ECL kit and the ChemiDoc Imaging System (Biorad).

### Fluorescence microscopy

Fluorescence images were acquired using a Leica SP8 confocal microscope equipped with a water immersion objective lens (× 25, numerical aperture 1.20; PL APO) (Imaging TRI-Genotoul plateform). GFP and RFP fluorescence was excited with the 488 nm ray line of the argon laser or the 561 nm ray line of the He-Ne laser respectively. The emission recording bands were set in the 505 to 530 nm range for GFP detection and 520 and 580 nm range for RFP detection. Image acquisition was done in the sequential mode using Leica LCS software and analyzed using the ImageJ software. Representative confocal images are shown after histogram normalization.

### Bimolecular fluorescence complementation (BiFC)

The coding sequences of *RSK1*, *MIK2* and *ZAR1* were amplified by PCR from Arabidopsis Col-0 cDNA using the primers indicated in Supplemental Table 4. To generate C-terminal fusion proteins with the C-terminal or N-terminal fragment of YFP, the genes were cloned into the expression vectors pEG100 (Early *et al*., 2006) using the Gateway technology (Invitrogen) (Gu and Innes 2011). The constructs expressing cYFP-or nYFP-tagged proteins in pEG100 were co-transfected into Arabidopsis Col-0 mesophyll protoplasts according to the previously described protocol (Yoo *et al*., 2007). The fluorescence was analyzed 16h after transfection by Confocal Laser Microscopy (Leica, SP8). Representative protoplasts were photographed with 510–550 nm (for YFP).

### Sub-network reconstruction

Interactors of MIK2 were recovered from RLK network described in Xi *et al*. (2019) and completed with Arabidospis BioGRID protein interaction datasets Version 4.0.189 (Oughtred *et al*., 2019). RLK annotation “ligand-perceiving” and “co-receptor” groups were recovered from Xi *et al*. (2019). Plant processes for each gene were verified with the current literature. Protein-Protein Interactions were plotted with Cytoscape software V3.7.2.

## Supporting information

Supplemental Table S1. Gene expression measurement by RT-qPCR, 6hours after infiltration by Xcc568 at 2.108 cfu/mL

## Acknowledgments

We thank Mehdi Khafif for technical assistance in Arabidopsis transient expression assays, Cécile Pouzet for cell imaging and Ullrich Dubiella and Marie Invernizzi for experimental assistance with plasmid constructs and diverse protein interaction tests. This work was supported by the French Agence Nationale de la Recherche – ANR grant (RIPOSTE ANR-14-OE19-0024-01) and the French Laboratory of Excellence Project TULIP (ANR-10-LABX-41).

F.D. is funded by a grant from Région Occitanie and the Plant Health & Environment Division of INRAE.

## Supplemental information

**Supplemental Figure S1.**
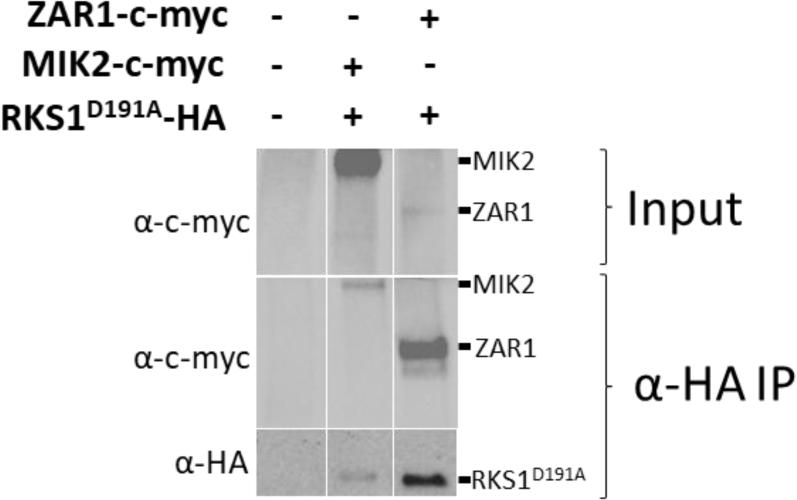
MIK2 full length interacts with RKS1. Co-Immunoprecipitation experiments with protein extracts from *N. benthamiana* inoculated by with the C58C1 *Agrobacterium* strain carrying *ZAR1-c-myc*, *MIK2-c-myc* constructs in one hand, and in the other hand, with a strain carrying *RKS1^D191A^-HA*. Input proteins were extracted and revealed by anti-HA and anti-c-myc HRP immunoblotting. Then, total proteins were subjected to anti-HA immunoprecipitation (α-HA IP), separated on a 4-15% SDS-PAGE gel and detected by anti-HA and anti-c-myc immunoblotting.

**Supplemental Figure S2.**
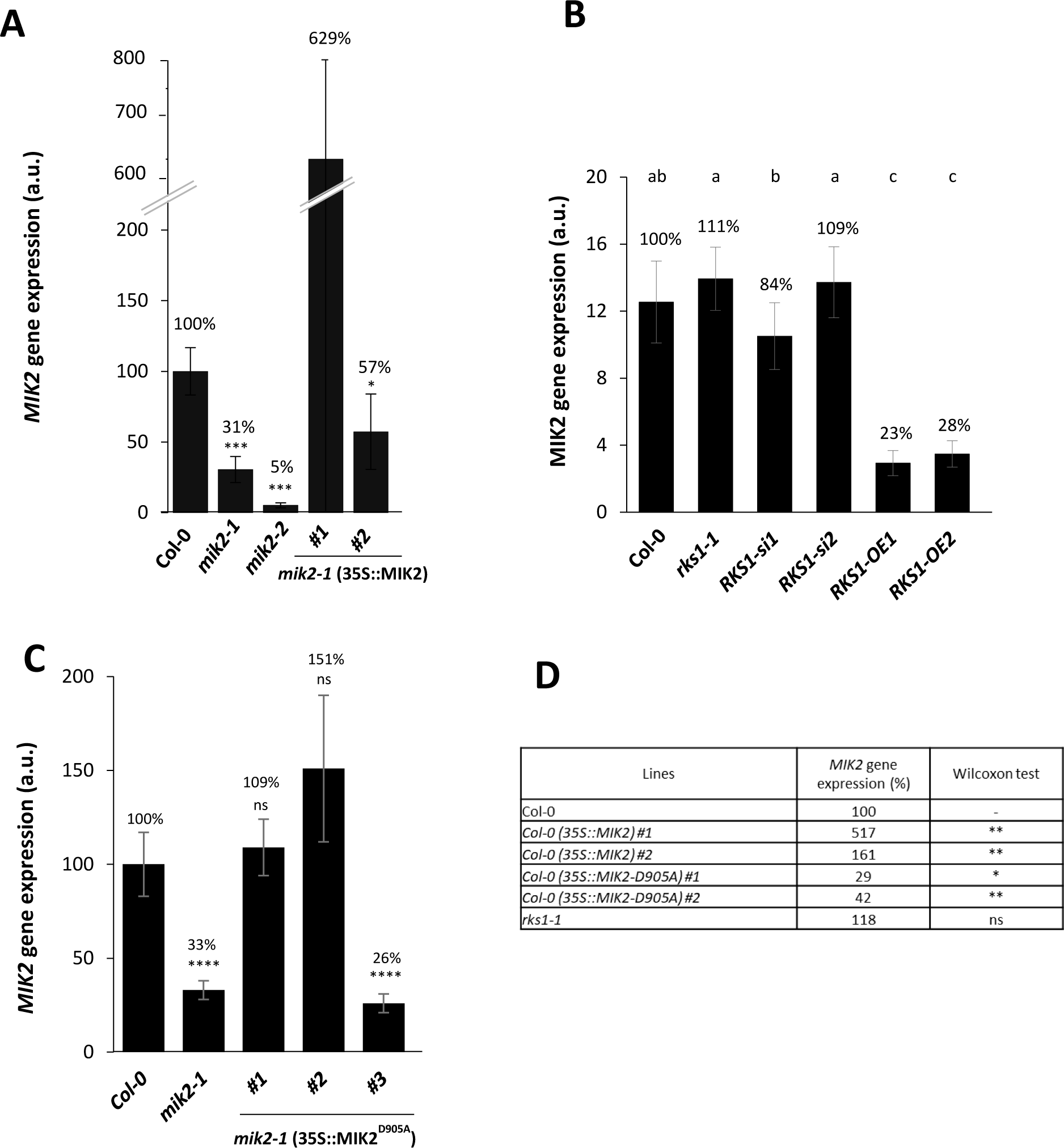
*MIK2* gene expression in mutants and other lines used in this study. **A,** *MIK2* gene expression was measured by RT-qPCR in healthy leaves in the mutants (*mik2-1* and *mik2-2)*, the *mik2-1* complemented lines by 35S::MIK2 (#1 and #2) and wild type plants (Col-0) type using primers after *mik2* T-DNA insertion. **B,** *MIK2* gene expression was measured by RT-qPCR in leaves, 6h after inoculation with a *Xcc568* bacterial suspension adjusted to 2.10^8^ cfu/mL in the wild type (Col-0), the mutant (*rks1-1)*, the *RKS1* amiRNA lines (*RNAi-si1 and RNAi-si2*), and in the *RKS1* overexpressor lines (*RKS1-OE1* and *RKS1-OE2).* Statistical groups were generated with the Wilcoxon test, based on 6 plants/line. **C,** *MIK2* gene expression was measured by RT-qPCR in healthy leaves in *mik2-1* mutant and *mik2-1* (35S::MIK2^D905A^) complemented lines and in the wild type using primers localized after *mik2-1* T-DNA insertion. Statistical analysis were performed with Wilcoxon test compared to Col-0. * indicate p-value < 0,05, ** indicate p-value < 0,01 and **** indicate p-value <0,0001. **D,** *MIK2* gene expression measured by RT-qPCR in lines overexpressing *MIK2* or MIK2^D905A^ in Col-0 background, in *rks1-1 mutant* and wild type plants. Statistical analyses were generated with the Wilcoxon test, based on 6 plants/line. *Represents *MIK2* gene expression significantly different than the wild type accession Col-0 (*pvalue ≤ 0.05 and ** pvalue ≤ 0.01, ns = not significantly different).

**Supplemental Figure S3.**
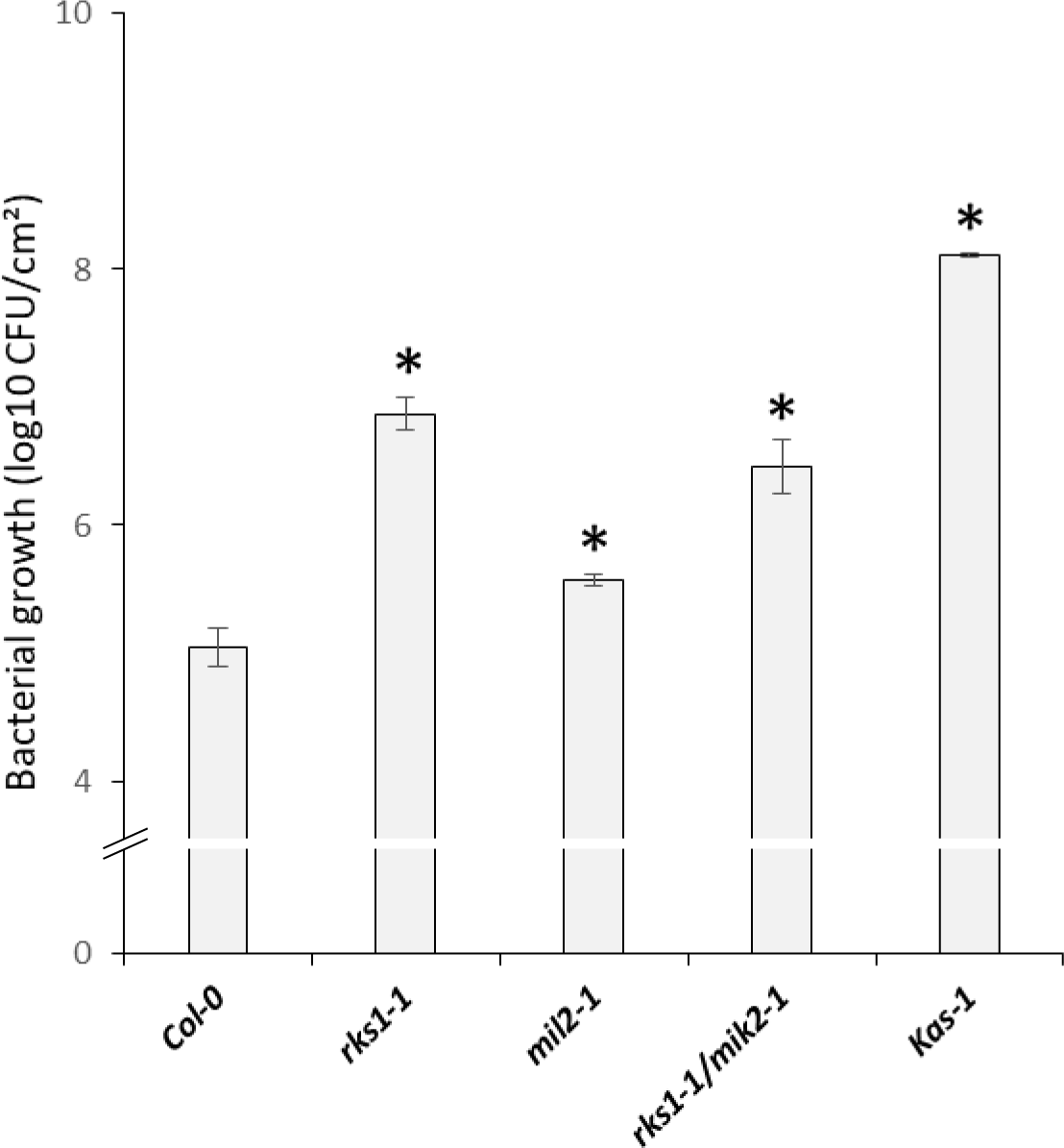
Analysis of *rks1-1*, *mik2-1* and *rks1-1/ mik2-1* mutant lines in response to *Xcc568*. Bacterial growth measurement (colony forming unit (CFU)/cm^2^ expressed in a log10 scale) in leaves of the wild type accession Col-0, the susceptible accession Kas-1 and *rks1-1*, *mik2-1* and *rks1-1/ mik2-1* mutant lines. Bacterial growth has been measured 7 days after inoculation with *Xcc568* at distance from the inoculation zone (at the tip of the inoculated leaves) with a bacterial suspension adjusted to 10^9^ CFU/mL. Data were collected from three independent experiments, each timepoint corresponds to measurements on 3–5 individual plants (four leaves/plant). Statistical analysis was performed using Kruskall-Wallis test and bacterial growth of Col-0 at day 7 as reference.

**Supplemental Figure S4.**
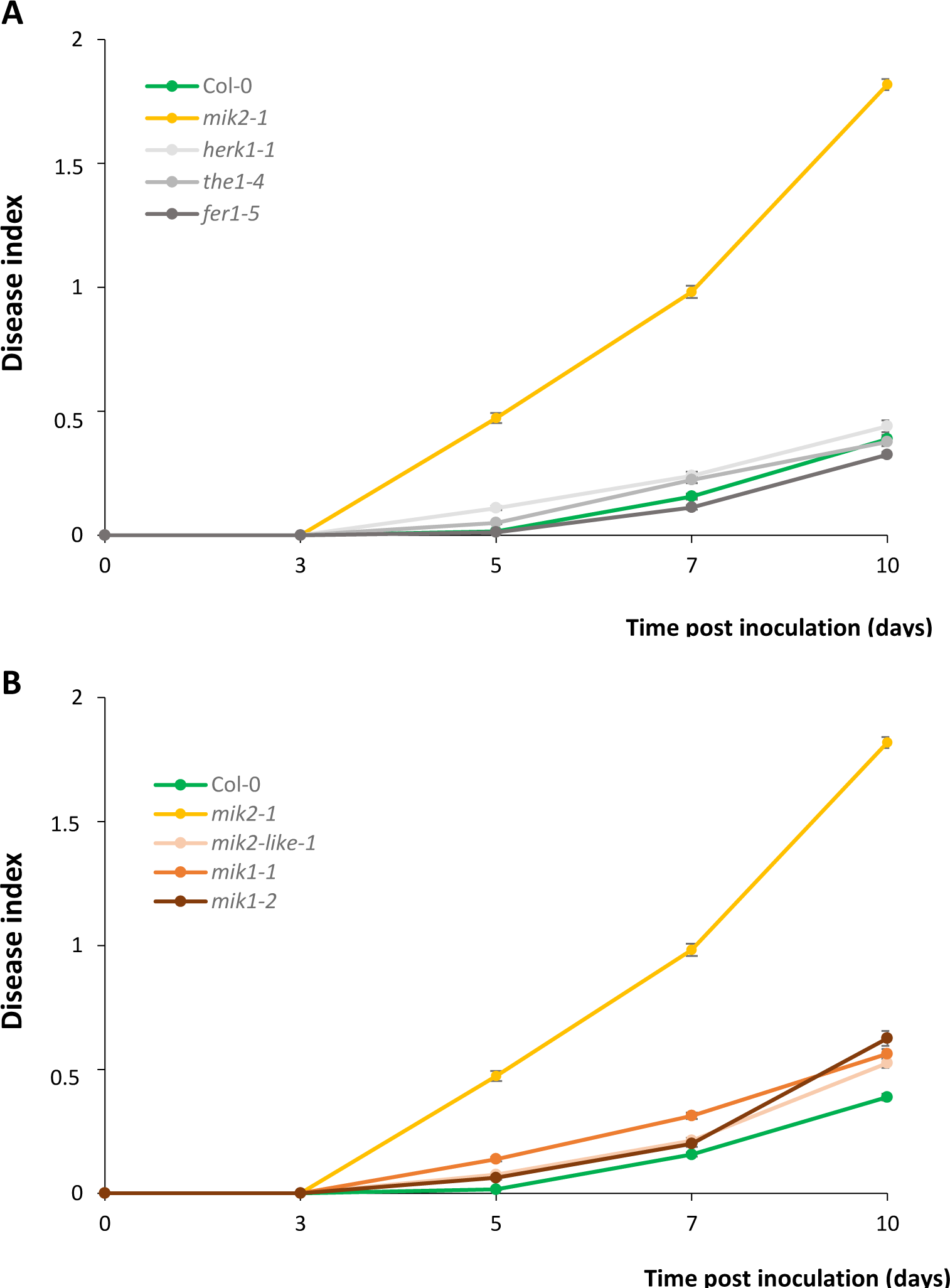
Cell wall sensing mutants are not affected in QDR to *Xcc568*. **A,** Time course evaluation of disease index after inoculation with a bacterial suspension adjusted to 2.10^8^ cfu/mL of *Xcc568* of Cell wall sensing mutants (*herk1-1, the1-4* and *fer1-5*) described in Van der Does *et al.,* 2017. **B,** Mutants affected in genes encoding MIK2 homologs (*mik2-like-1*, *mik1-1* and *mik1-2*). Means and standard errors were calculated from 5 plants per line and based on 3 independent experiments. * Represents comparison to Col-0 based on comparison of disease index kinetics.

**Supplemental Table S1.** Gene expression measurement by RT-qPCR, 6hours after infiltration by *Xcc568* at 2.10^8^ cfu/mL. (Deposited online)

**Supplemental Table S2.**
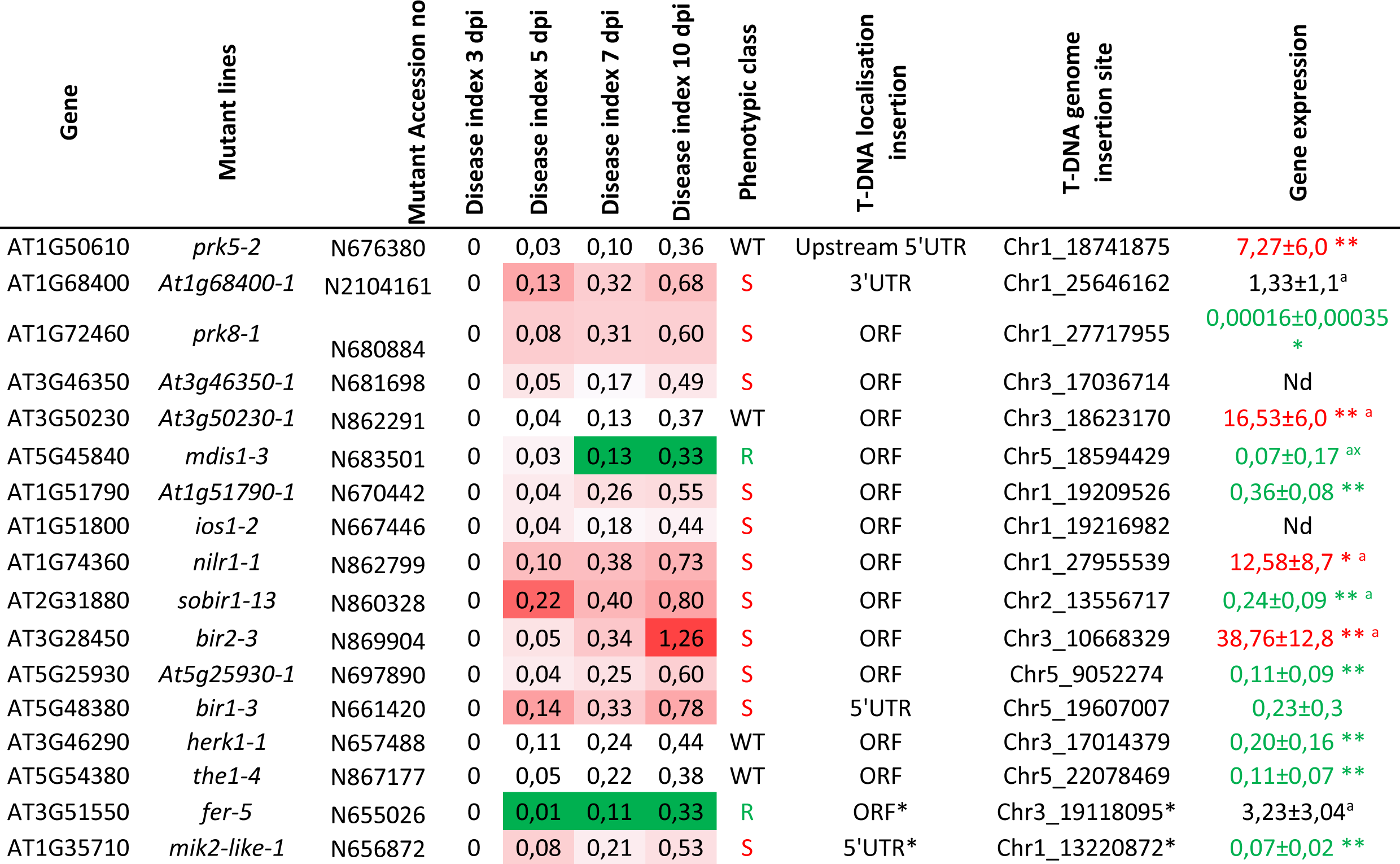
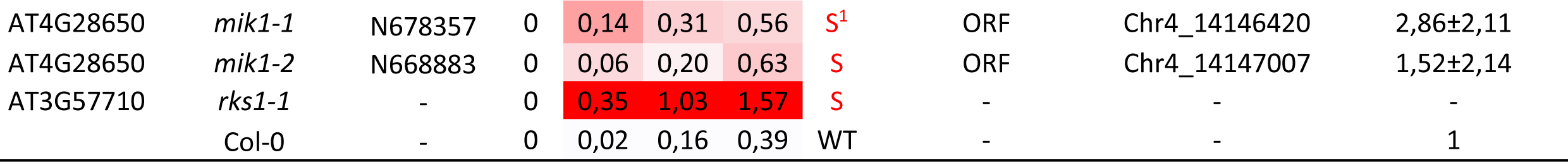
Molecular analysis of insertional mutants for RLK network. Mutant lines, corresponding gene accessions, mutant phenotypes after inoculation with a bacterial suspension of Xcc568 adjusted to 2.108 cfu/mL and gene expression in corresponding mutants. A heatmap highlights time course evaluation of mutant disease index as compared to the wild type (Col-0) disease index. Disease symptoms were observed on leaves of mutant and wild-type plants at 3-, 5-, 7- and 10-days post-inoculation (dpi). Means were calculated from 20-25 plants on 4 to 5 experiments. Green represents disease index significantly reduced as compared to Col-0 (more resistant), Red represents disease index significantly increased as compared to Col-0 (more susceptible) and white, not significantly different from Col-0. Based on kinetic modeling deference with Col-0, the phenotype is indicated as R (resistant), S (susceptible) or WT (not significantly different from Col-0). 1 indicate mutant seems more susceptible but not significantly from Col-0. For each mutant, location of the T-DNA insertion was determined by sequencing and indicated (5’UTR, ORF or 3’UTR regions), * indicate predicted location of the T-DNA from TAIR. Gene expression was evaluated by Q-RT-PCR in the mutant plants, and indicated in red (increased in the mutant as compared to Col-0), green (decreased in the mutant as compared to Col-0) and in black (not affected in the mutant as compared to Col-0) in the last column. Statistical analysis was performed by comparing the kinematic of the average disease index of each mutant to the kinematic of the average disease index of Col-0 (see Material and Methods section). * and ** represent significate differences generated with Wilcoxon test, * = P-value <0,05, **=P-value <0,001. Q-RT-PCR was performed with primers downstream the T-DNA insertion except for mutants mentioned by a.

**Supplemental Table S3.** Molecular analysis of insertional mutants for RKS1 dependent subnetwork. Mutant lines, corresponding gene accessions, mutant phenotypes after inoculation with a bacterial suspension of Xcc568 adjusted to 2.108 cfu/mL and gene expression in corresponding mutants. A heatmap highlights time course evaluation of mutant disease index as compared to the wild type (Col-0) disease index. Disease symptoms were observed on leaves of mutant and wild-type plants at 3-, 5-, 7- and 10-days post-inoculation (dpi). Means were calculated from 15-25 plants on 3 to 5 experiments. Red represents disease index significantly increased as compared to Col-0 (more susceptible). Based on kinetic modeling deference with Col-0, the phenotype is indicated S (susceptible) or WT (not significantly different from Col-0). For each mutant, location of the T-DNA insertion was determined by sequencing and indicated (5’UTR, ORF or 3’UTR regions). Gene expression was evaluated by Q-RT-PCR in the mutant plants, and indicated in green (decreased in the mutant as compared to Col-0) a in the last column. Statistical analysis was performed by comparing the kinematic of the average disease index of each mutant to the kinematic of the average disease index of Col-0 (see Material and Methods section). * Represents significate differences generated with Wilcoxon test, * = P-value <0,05. Q-RT-PCR was performed with primers downstream the T-DNA insertion.

| Gene | Mutant lines | Mutant Accession no | Disease index 3 dpi | Disease index 5 dpi | Disease index 7 dpi | Disease index 10 dpi | Phenotypic class | T-DNA localisation insertion | T-DNA genome insertion site | Gene expression |
| --- | --- | --- | --- | --- | --- | --- | --- | --- | --- | --- |
| AT4G33430 | *bak1-3* | N662069 | 0 | 0,16 | 0,61 | 1,04 | S | ORF | Chr4_16089269 | 0,2±0,03 * |
| AT4G33430 | *bak1-4* | N663745 | 0 | 0,24 | 0,59 | 1,29 | S | ORF | Chr4_16087795 | 0,007±0,001* |
| AT5G20480 | *efr-1* | N799998 | 0 | 0,04 | 0,34 | 0,86 | S | ORF | Chr5_6924546 | 0,4±0,02 |
| AT5G46330 | *fls2-3* | N668000 | 0 | 0,09 | 0,45 | 0,88 | S | ORF | Chr5_18794872 | 0,1±0,2 * |
|  | *rks1-1* |  | 0 | 0,50 | 1,30 | 1,86 | S | - | - | - |
|  | Col-0 |  | 0 | 0,05 | 0,28 | 0,55 | WT | - | - | 1 |

**Supplemental Table S4.**
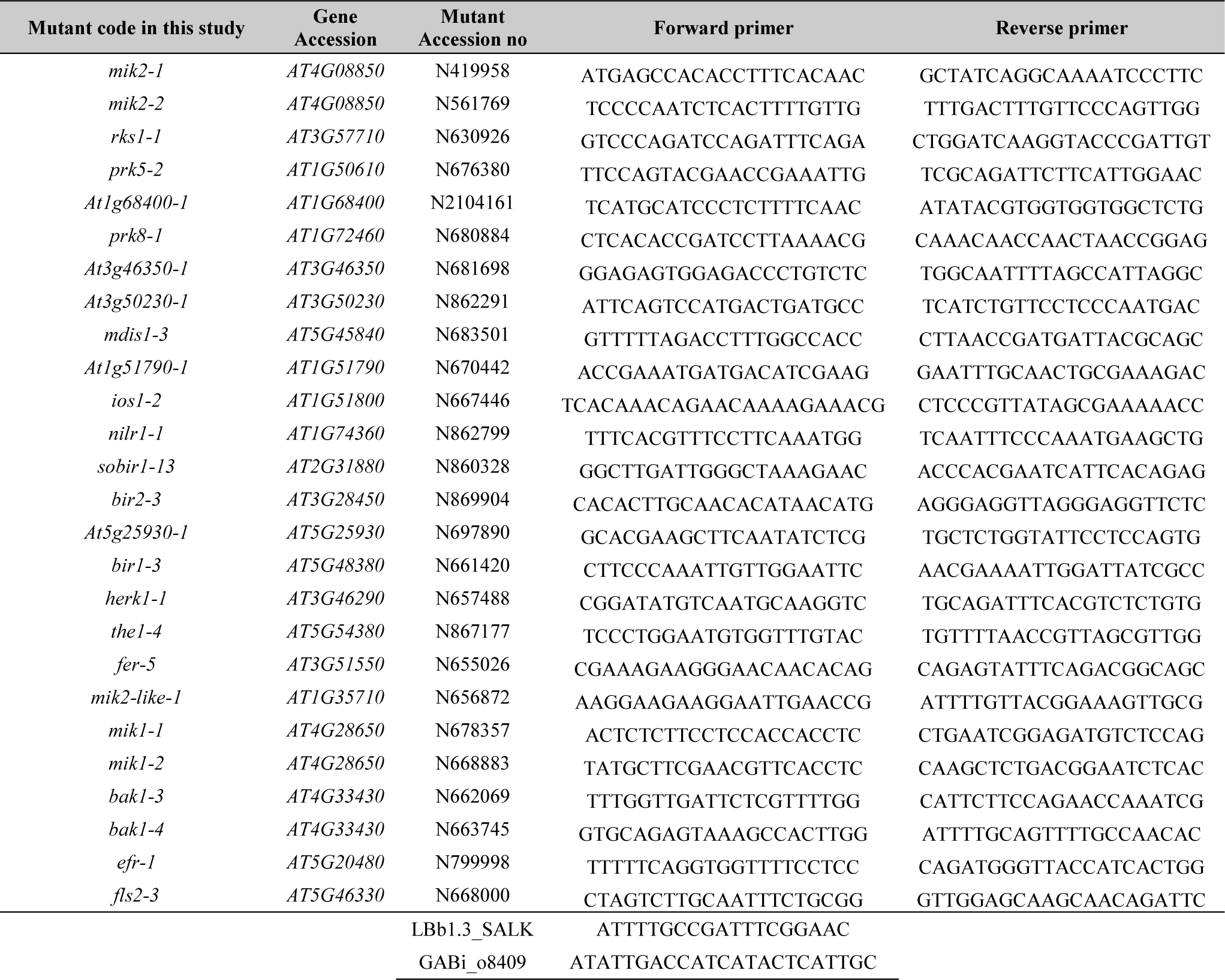
List of primers and oligonucleotide sequences used for mutant genotyping.

**Supplemental Table S5.**
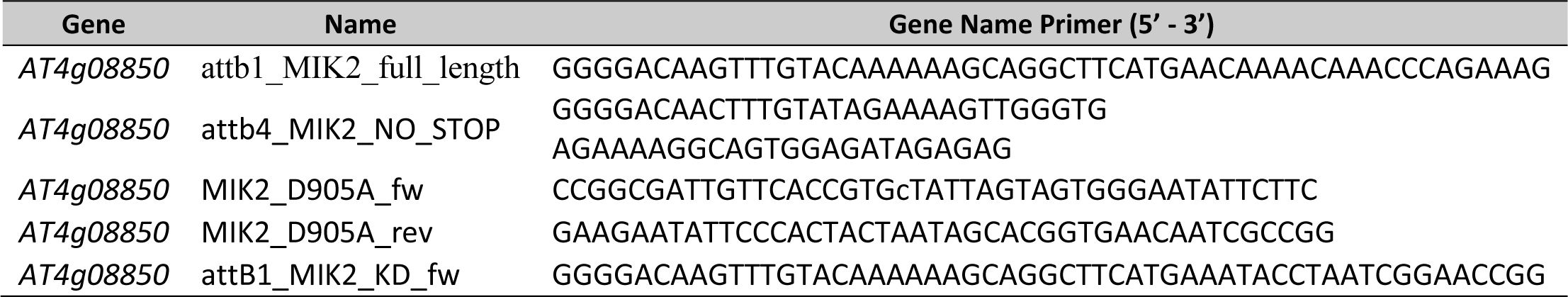
Primers used for MIK2 constructions.

**Supplemental Table S6.** List of plasmid constructs in pEarlyGate100 used in this study.

| Gene | Gene construction | C-terminal tags |
| --- | --- | --- |
| *AT4G08850* | MIK2 | 3xHA |
| *AT4G08850* | MIK2 | 5xc-myc |
| *AT4G08850* | MIK2 | mRFP1 |
| *AT4G08850* | MIK2<sup>D905A</sup> | 3xHA |
| *AT4G08850* | MIK2<sup>D905A</sup> | mRFP1 |
| *AT4G08850* | MIK2_KinaseDomain | 3xHA |
| *AT4G08850* | MIK2<sup>D905A</sup>_KinaseDomain | 3xHA |
| *AT3G57710* | RKS1 | 5xc-myc |
| *AT3G57710* | RKS1 | eGFP |
| *AT3G57710* | RKS1<sup>D191A</sup> | 5xc-myc |
| *AT3G57710* | RKS1<sup>D191A</sup> | eGFP |
| *AT3G50950* | ZAR1 | 5xc-myc |

**Supplemental Table S7.**
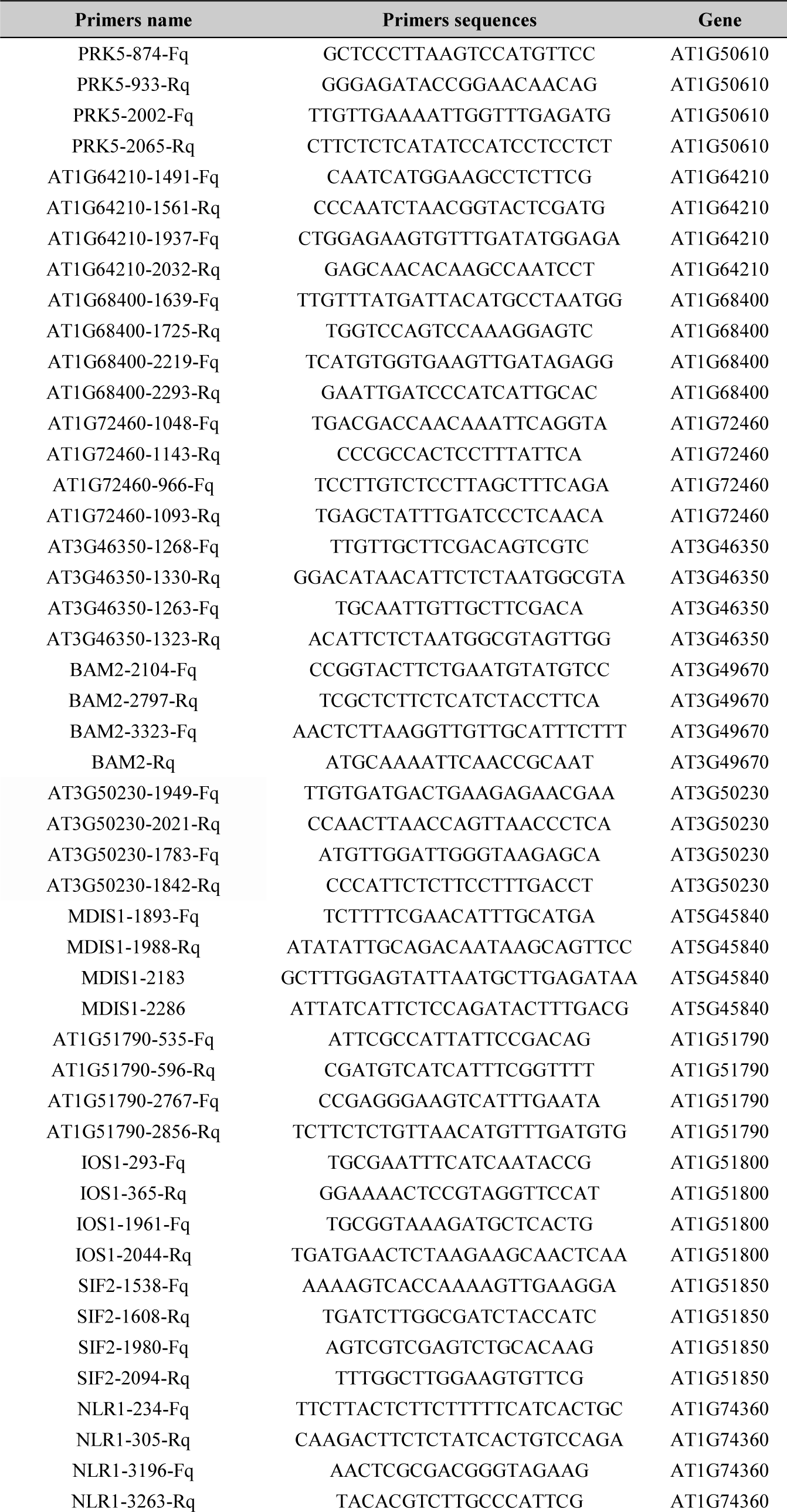
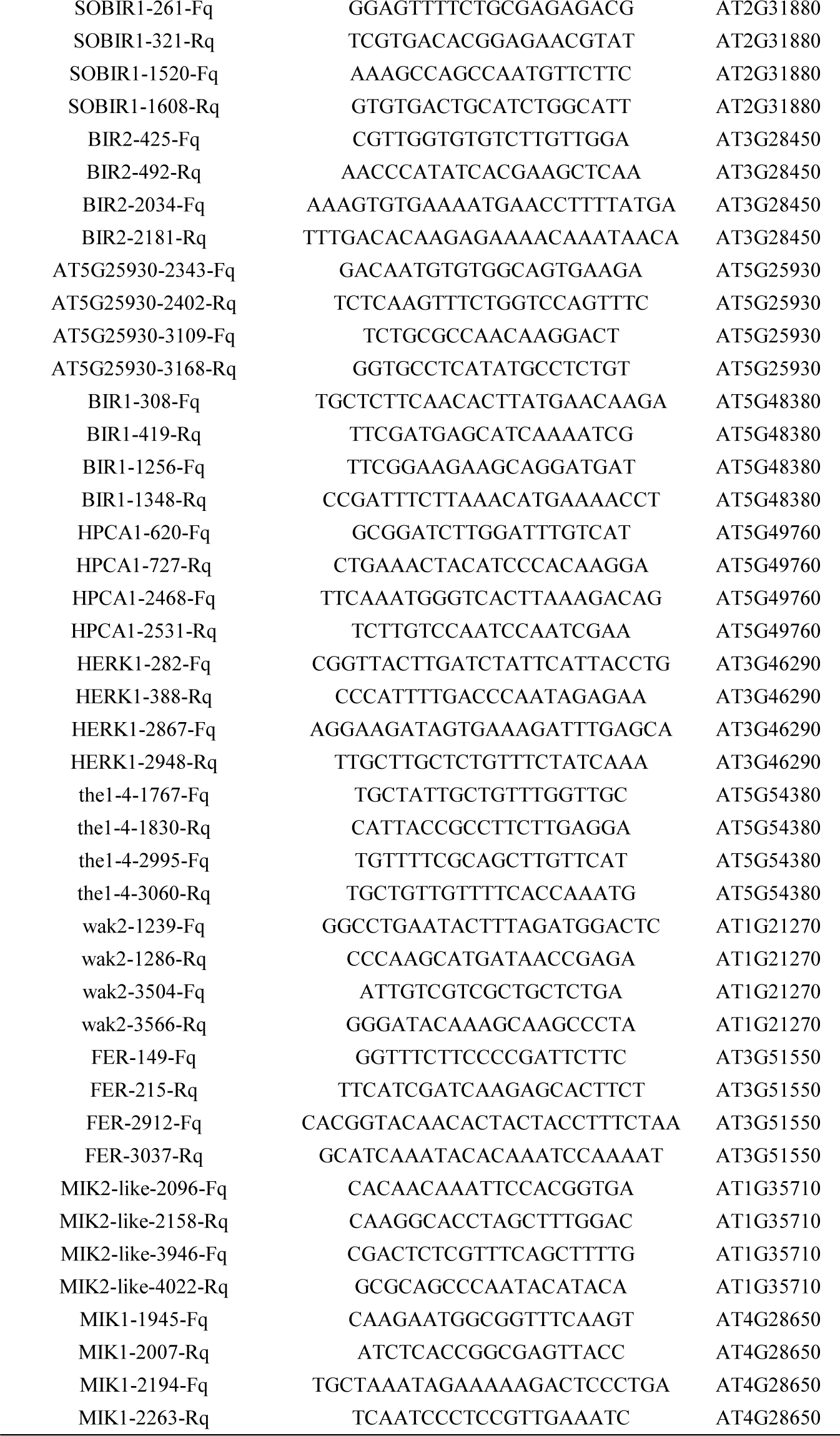
List of primers used for RT-qPCR for expression profile evaluation.

**Supplemental Table S8.**
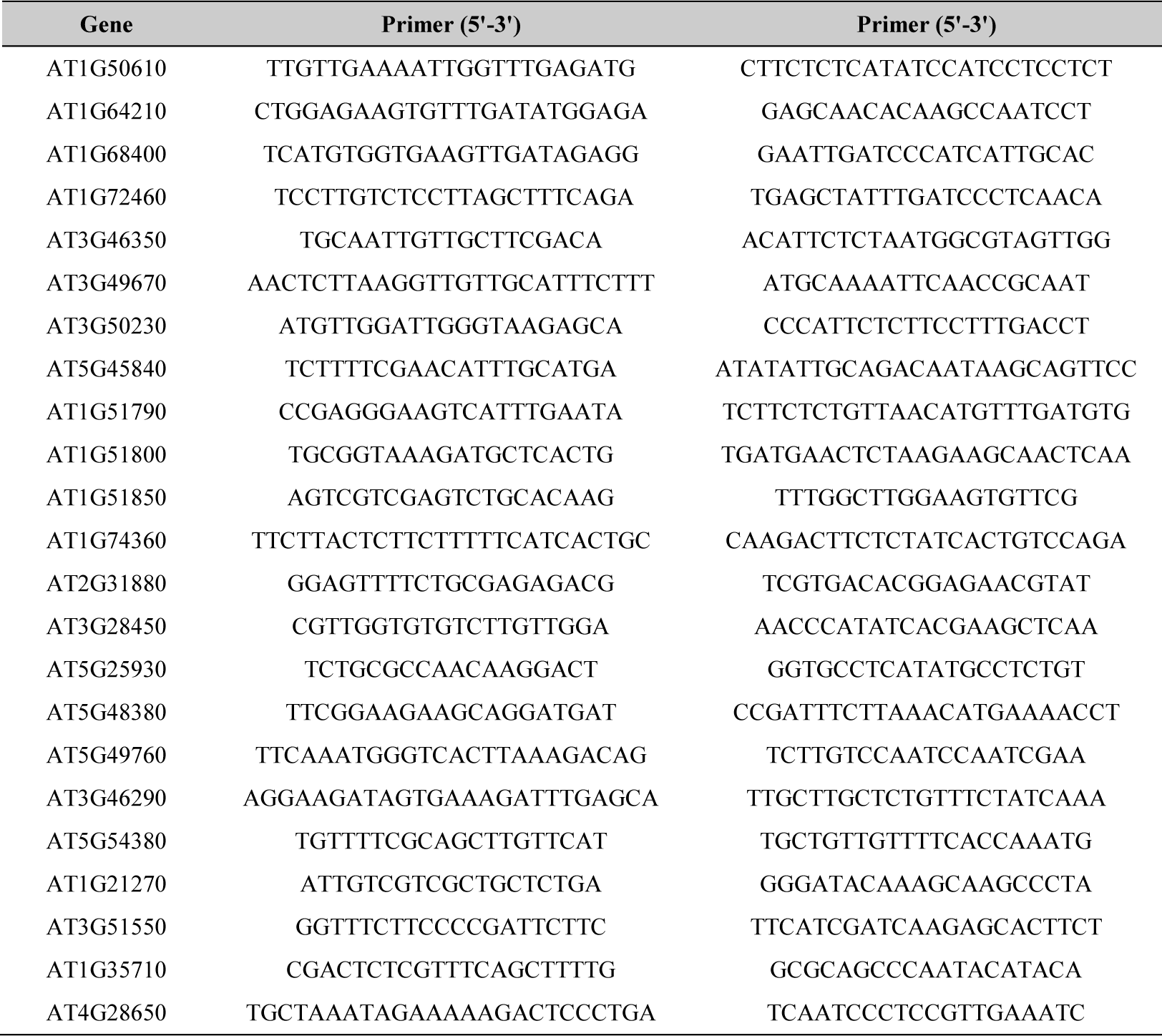
List of primers used for RT-qPCR molecular characterization of T-DNA mutants.

**Dataset 1.** Protein-protein interactions used for the generation of the RKS1/MIK2 PPI network.

| Node 1 | Node 2 | Source |
| --- | --- | --- |
| AT1G64210 | BAM2 | Biogrid-db |
| AT1G64210 | FLS2 | Biogrid-db |
| AT1G68400 | BAM1 | Biogrid-db |
| AT1G68400 | BAM2 | Biogrid-db |
| AT1G72460 | SOBIR1 | Biogrid-db |
| AT1G72460 | BAM1 | Biogrid-db |
| AT3G50230 | ERL2 | Biogrid-db |
| AT5G25930 | BAM1 | Biogrid-db |
| AT5G25930 | BAM2 | Biogrid-db |
| BAK1 | EFR | Biogrid-db |
| BAM1 | BRI1 | Biogrid-db |
| BAM1 | EFR | Biogrid-db |
| BIR1 | BAK1 | Biogrid-db |
| BIR1 | BIR1 | Biogrid-db |
| BIR1 | FLS2 | Biogrid-db |
| BIR2 | BAK1 | Biogrid-db |
| BRI1 | BSK3 | Biogrid-db |
| BRI1 | EFR | Biogrid-db |
| BSK3 | BSK3 | Biogrid-db |
| ERL2 | EFR | Biogrid-db |
| ERL2 | ERL2 | Biogrid-db |
| IOS1 | BSK3 | Biogrid-db |
| MDSI1 | SOBIR1 | Biogrid-db |
| MIK2 | AT1G64210 | Biogrid-db |
| MIK2 | AT1G68400 | Biogrid-db |
| MIK2 | AT1G72460 | Biogrid-db |
| MIK2 | AT3G50230 | Biogrid-db |
| MIK2 | MDSI1 | Biogrid-db |
| MIK2 | PRK4 | Biogrid-db |
| MIK2 | AT1G49100 | Biogrid-db |
| MIK2 | AT3G46350 | Biogrid-db |
| MIK2 | BAM2 | Biogrid-db |
| MIK2 | BSK3 | Biogrid-db |
| MIK2 | PRK5 | Biogrid-db |
| MIK2 | RKS1 | Biogrid-db |
| PRK4 | BAM1 | Biogrid-db |
| PRK5 | EFR | Biogrid-db |
| RKS1 | EFR | Biogrid-db |
| SIF2 | BAK1 | Biogrid-db |
| SOBIR1 | BAK1 | Biogrid-db |
| SOBIR1 | BAM1 | Biogrid-db |
| EFR | FLS2 | Makowska-Luzan *et al.*, 2018 doi:10.1038/nature25184 |
| MIK2 | AT1G51790 | Wang, T., et al. (2016). doi: 10.1038/nature16975. |
| MIK2 | AT5G25930 | Xi *et al.*, 2019 doi:10.3389/fpls.2019.00472 |
| MIK2 | BIR2 | Xi *et al.*, 2019 doi:10.3389/fpls.2019.00472 |
| MIK2 | SIF2 | Xi *et al.*, 2019 doi:10.3389/fpls.2019.00472 |
| MIK2 | SOBIR1 | Xi *et al.*, 2019 doi:10.3389/fpls.2019.00472 |
| MIK2 | BIR1 | Xi *et al.*, 2019 doi:10.3389/fpls.2019.00472 |
| MIK2 | HPCA1 | Xi *et al.*, 2019 doi:10.3389/fpls.2019.00472 |
| MIK2 | IOS1 | Xi *et al.*, 2019 doi:10.3389/fpls.2019.00472 |
| MIK2 | NIR1 | Xi *et al.*, 2019 doi:10.3389/fpls.2019.00472 |
| IOS1 | BAK1 | Yeh *et al.*, 2016 doi:10.1105/tpc.16.00313 |
| IOS1 | EFR | Yeh *et al.*, 2016 doi:10.1105/tpc.16.00313 |
| IOS1 | FLS2 | Yeh *et al.*, 2016 doi:10.1105/tpc.16.00313 |
| MIK2 | BAK1 | Rhodes *et al.*, 2021 doi.org/10.1038/s41467-021-20932-y |

## Notes

### Competing Interest Statement

The authors have declared no competing interest.

